# Structure of the Pre-Initiation Complex Explains CMGE Biogenesis

**DOI:** 10.64898/2026.05.15.725318

**Authors:** Thomas Pühringer, Berta Canal, Giacomo Palm, Agata Butryn, Emma C. Couves, Oliver Willhoft, Jacob S. Lewis, John F. X. Diffley, Alessandro Costa

## Abstract

When cells enter S phase, bidirectional DNA replication is initiated through the kinase-regulated recruitment of three activators (Cdc45, GINS and Pol epsilon) to a duplex DNA-loaded double hexamer of MCM ATPases. Together these proteins form two CMGE helicases that establish divergent replication forks as they become separated^1^. To understand CMGE biogenesis, we reconstituted the pre-Initiation Complex with purified yeast proteins. The cryo-EM structure shows a set of firing factors caught in the act of assembling two symmetric CMGEs. We show how stepwise complex formation reshapes MCM in preparation for DNA opening and we explain how ATP promotes firing-factor ejection and CMGE maturation. While we find that Sld2 promotes GINS recruitment to MCM as expected, it also aids efficient separation of the CMGE dimer, and it is essential for lagging strand ejection from MCM. These findings have direct implications for our understanding of the metazoan Sld2 ortholog, RECQL4, pointing to a replication-fork establishment mechanism conserved across eukaryotes.

## Introduction

DNA replication must occur only once per cell cycle to maintain genome stability. To achieve this, eukaryotes have evolved to temporally separate the loading of the replicative helicase from its activation^1^. During the G1 phase of the cell cycle, two copies of the Minichromosome maintenance (MCM) motor of the replicative helicase are loaded as an inactive double hexamer (DH) onto DNA replication origins^2,3^. Activation occurs upon S-phase transition and involves the recruitment of Cdc45 and the tetrameric GINS (Go-Ichi-Ni-San) complex to MCM, together forming the Cdc45-MCM-GINS (CMG) helicase^4,5^. CMG assembly occurs under the control of three kinases. It is promoted by the Dbf4-dependent kinase (DDK)^6-8^ and Cdc28-Clb5 (hereafter, CDK), whose activity increases when cells enter S phase^9-11^. CMG assembly is instead inhibited by the checkpoint kinase Rad53 that blocks late origin firing if DNA damage is detected^9-13^. DDK selectively phosphorylates DNA-loaded DHs^14^. A heterodimeric firing factor, composed of Sld3 (essential) and Sld7 (dispensable), then recognises the phosphorylated DH and recruits Cdc45 to MCM^10,11,15^. An N-terminal truncation of Mcm4 bypasses the requirement for DDK in cells^7,8^ and cryo-electron microscopy (cryo-EM) work showed that phosphorylation causes N-terminal Mcm4 to become unstructured^16,17^. Whether Sld3 engages an epitope unmasked upon N-terminal Mcm4 phosphorylation or merely reads phospho-Mcm4 sites remains to be established. Also, while we know that Rad53 prevents CMG formation by targeting Sld3 (alongside DDK^13^), the mechanism is unclear. The second activating kinase, CDK, targets two firing factors, Sld3 and Sld2, previously implicated in GINS recruitment. Phospho-Sld2 and phospho-Sld3 are recognised by the Dpb11 phospho-reader^10,11^. The leading strand polymerase Pol epsilon (formed of Pol2, Dpb2, Dpb3 and Dpb4) also contributes to CMG assembly and becomes part of the holo-helicase, CMGE. In particular, the N-terminal domain of Dpb2 supports CMG formation in cells^18^ and a complex containing Dpb2 and the C-terminal half of Pol2 achieves GINS recruitment and replication initiation reconstituted with purified proteins^19^.

In the inactive DH, an Mcm7-specific N-terminal insertion (NTI) from one MCM hexamer reaches across the DH interface and protects the N-terminal A domain of Mcm5 of the opposed hexamer^20^ (**Fig. 1a**). The same Mcm5 A domain site is engaged by GINS in the CMG, implying that Mcm7 must let go of Mcm5 for GINS to bind^21,22^. How this happens is unknown. Likewise, it is established that Cdc45 and GINS recruitment occur sequentially^23-25^. Whether recruitment of each component involves only one or both MCM hexamers at once is debated^25-27^.

**Figure 1.**
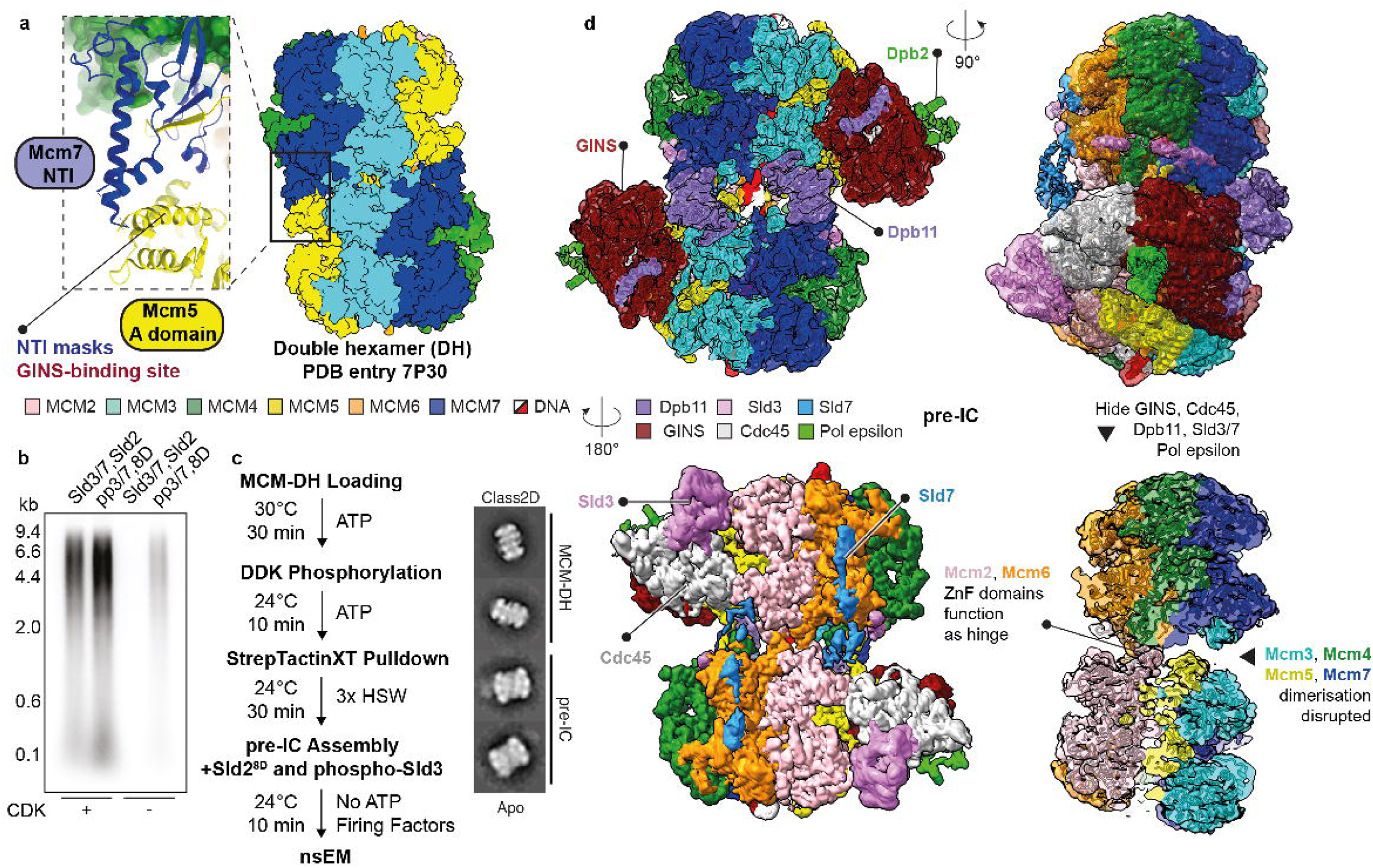
Double hexamer and pre-IC structures. **(a)** In the DH, the Mcm7 NTI element from one MCM ring protects the N-terminal A domain of the opposed ring. **(b)** CDK-pre-phosphorylated Sld3/7 and the Sld2 8D phospho-mimetic mutant support origin-dependent replication reconstituted in a test tube, albeit at lower level (This experiment was performed two times). The low levels could be due to either the Sld2-8D phosphomimetic mutations only covering a subset of CDK sites, or to inefficient phosphorylation of Sld3 in the isolated pre-phosphorylation reaction, compared to the complete initiation mix. For gel source data, see Supplementary Fig. 1. **(c)** CMG assembly in the absence of ATP, using CDK-pre-phosphorylated Sld3/7 and Sld2 8D enables pre-initiation complex formation. **(d)** Cryo-EM structure of the pre-IC. The two MCMs are split on one side, with Mcm2-6 functioning as a hinge.

Origin activation is controlled by ATP binding and hydrolysis. MCM loading, for example, requires ATP hydrolysis^28,29^, so that, by the time a DH is formed, eight of its twelve subunits are bound to ADP^14^. Firing factor recruitment to the DH promotes ADP release, and ATP binding by MCM achieves stable double CMG-Pol epsilon (ATP-dCMGE) complex formation, which nucleates DNA melting^24,30^. It has been postulated that a pre-Initiation Complex (pre-IC) might exist^31^, where Sld2, Sld3/7 and Dpb11 all bind the DH at the same time, while also recruiting Cdc45, GINS and Pol epsilon^24,32^. However, neither has the pre-IC ever been isolated, nor do we know how Sld2, Sld3/7 and Dpb11 are ejected, achieving ATP-dCMGE maturation^24,30^. Finally, whether the role of Sld2 is conserved across eukaryotes is unclear. In fact, yeast Sld2 functions in CMGE assembly in yeast, while the metazoan ortholog RecQL4 has been implicated in a loosely defined downstream activation step. To address these issues and understand replisome biogenesis, we have taken a biochemical reconstitution approach, combined with cryo-EM imaging and single-particle reconstruction.

## Results

### Cryo-EM structure of the pre-IC

In earlier cryo-EM work we established that, by the time the ATP-dCMGE complex is formed at an origin, all factors involved in its assembly process have been released^30^. We reasoned that, should ATP binding by MCM promote this release, forming dCMGE in a buffer that lacks ATP may retain firing factors Sld2, Sld3/7 and Dpb11 at origins, allowing us to reconstitute the pre-IC. Obtaining enough sample for cryo-EM analysis in such conditions represents a challenge. In fact, ATP is required for DH phosphorylation by DDK^33^ as well as Sld2 and Sld3 phosphorylation by CDK^10,11^. To circumvent this issue, we isolated the phospho-DH on origin DNA^30^, in a buffer lacking ATP (**Supplementary Figs. 2a-e**). Inspired by earlier work^24^, we also phosphorylated recombinant Sld3/7 using CDK and re-purified it in ATP-free conditions (pre-phosphorylated Sld3/7, hereafter ppSld3/7, **Supplementary Fig. 2f**). We also cloned, expressed and purified a phosphomimetic variant of Sld2 containing 8 of 11 aspartate substitutions (hereafter Sld2-8D), previously shown to support Dpb11 binding and CDK bypass in cells^10,34^ (**Supplementary Fig. 2g**). We then tested ppSld3/7 and Sld2-8D in a DNA replication reaction reconstituted in vitro with purified proteins^19^. DNA replication could be established in the absence of CDK with these reagents, though at reduced levels compared to reactions containing CDK (**Fig. 1b**). Despite the reduction, ATP-dCMGE complexes^30^ could be assembled efficiently using ARS1-loaded phospho-DH, ppSld3/7 and Sld2-8D in the absence of CDK, as observed by negative stain electron microscopy (nsEM, **Supplementary Figs. 3a,b**). We then asked whether GINS and Cdc45 could be recruited to MCM using ppSld3/7 and Sld2-8D, in the absence of CDK and any nucleotide. nsEM two-dimensional (2D) averaging revealed a complex reminiscent of ATP-dCMGE^30^, however with MCMs engaged in tighter interaction and no recognisable Pol epsilon density, at least at the limited resolution achieved with negative staining (**Fig. 1c**). Unlike ATP-dCMGE, the new ATP-free complex became disassembled when purified in high-salt conditions (**Supplementary Figs. 3c-e**). This observation agrees with earlier Western blot evidence that high-salt resistant association of GINS and Cdc45 to licensed-origin DNA requires ATP binding^24^. To determine the composition of the ATP-free high-salt sensitive assembly, we solved the cryo-EM structure (**Extended Data Figs.1,2**). Inspection of the resulting density map, refined to 3.4 Å resolution after symmetry expansion^35^ (or 3.2 Å for the locally-refined asymmetric unit), revealed a quasi-symmetric (flexible) assembly of two CMGs (**Extended Data Table 1**). Compared to the DH, Mcm5-3-7-4 are disengaged across the two rings. Homo-dimerisation via Mcm2 and Mcm6 persists as observed in the DH^20^, with Zinc Finger (ZnF) domains acting as a hinge. Unoccupied density could be assigned to two copies of Sld3, Sld7, Dpb11 and the Dpb2 subunit of Pol epsilon, which together with two CMG assemblies form the pre-IC complex (**Fig. 1d**).

### Kinase-regulated Mcm4 engagement by Sld3

Previous studies have established that DDK docking onto the DH dislodges the N-terminal tail of Mcm4 from the A domain of Mcm4 (A4, **Figs. 2a,b**)^16,17,36^. A4 remains occupied by the Dbf4 subunit of DDK, while the Cdc7 kinase subunit phosphorylates the N-terminal Mcm4 tail that has become solvent-exposed (**Figs. 2a,c**). When DDK releases the DH, the negatively charged phospho-Mcm4 tail does not return to binding the electronegative A4 site, probably due to electrostatic repulsion (**Figs. 2a,d**)^16,17^. Within the pre-IC, we find that the newly exposed A4 epitope becomes engaged by a C-terminal Sld3 alpha helix, which in turn extends towards the two neighbouring subunits on either side of Mcm4. This same interaction was also predicted using AlphaFold 3 (AF3)^37^. N-terminally, Sld3 touches the Mcm6 A domain and C-terminally it engages the Mcm7 NTI element (**Fig. 2e**). This Sld3 interaction may contribute to dislodging Mcm7 from its position across the two hexamers, where it protected the Mcm5 A domain in *trans*. We expressed and purified a truncated Sld3/7 complex lacking the Mcm4 binding site of Sld3 (Δ515-538), to test whether this variant can still form pre-IC, and support DNA replication in reconstituted reactions. We found that, compared to the wild-type protein, both functions are almost completely blocked when using the truncated variant (**Figs. 2f,g**). Our results agree with previous biochemical data, which identified the same Sld3 region as essential for MCM binding^23^. In addition to the interaction with Mcm4 described above, Sld3 has also been observed to bind specific phosphorylated segments of Mcm4 and Mcm6 independently (not observed in our structure). This indicates a distinct phospho-reader role for Sld3, which complements its function in recognising a DDK-dependent structural change in MCM^23^.

**Figure 2.**
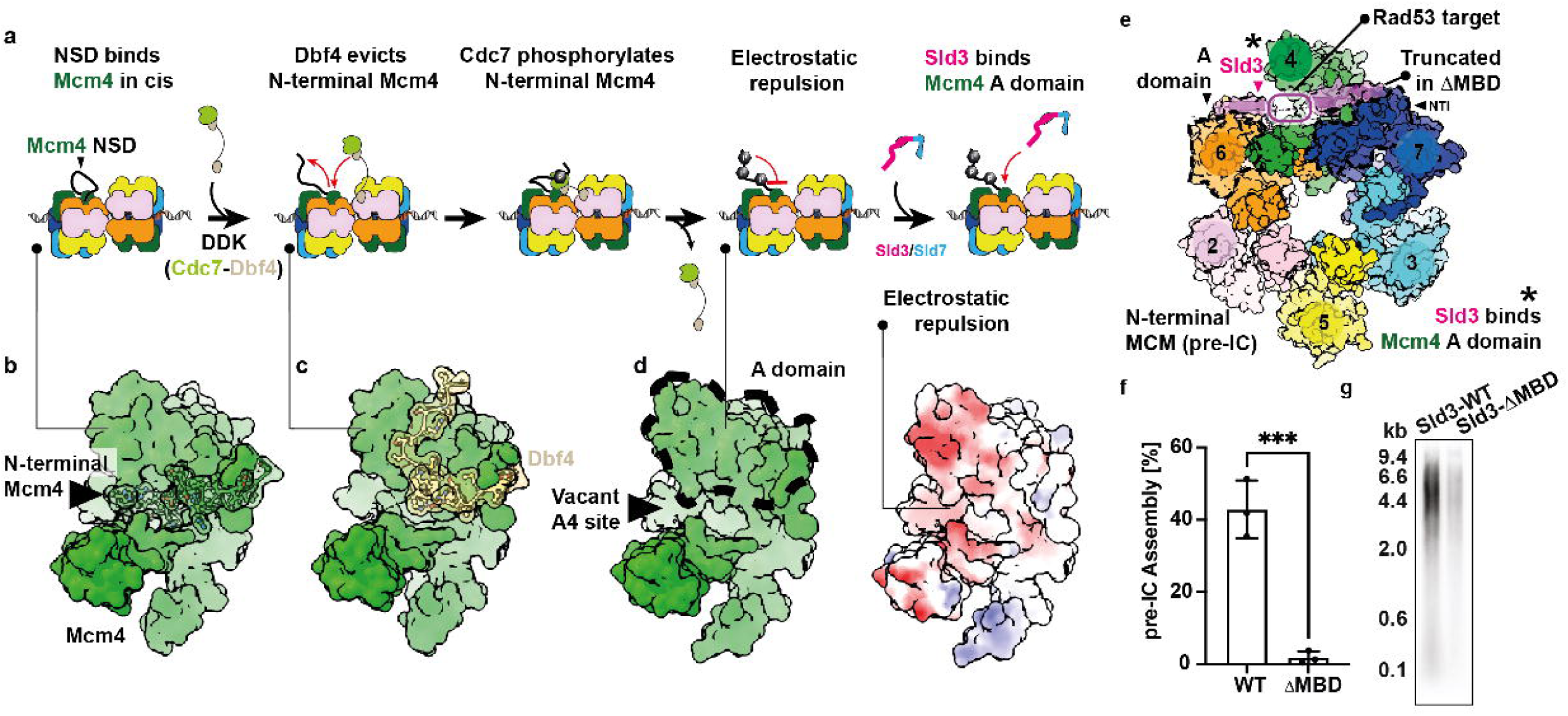
DDK enables Sld3 binding to Mcm4. **(a)** Cartoon depicting the sequence of events leading to Sld3 recruitment. First, DDK subunit, Dpb4, dislodges N-terminal Mcm4 from Mcm4 A domain. Then DDK phosphorylation of N-terminal Mcm4 prevents re-engagement to A domain. Finally, Sld3 engages newly exposed Mcm4 A domain epitope (vacant A4 site). **(b)** Structure of non-phosphorylated Mcm4 (PDB entry 7V3U). **(c)** Structure of Mcm4 engaged by Dbf4 subunit of DDK (PDB entry 7PT7). **(d)** Structure of phosphorylated Mcm4 (7P30). Surface representation of phosphorylated Mcm4, coloured by electrostatic potential. **(e)** Sld3 binds Mcm4 A domain as well as the neighbouring NTI Mcm7 element and Mcm6 A domain. **(f)** Truncation of the MCM binding domain of Sld3 impairs pre-IC assembly (P-value determined by two-tailed t-test; *\*\*\*P*=0.0010; Error bars, mean ± s.d. This experiment was performed three times) and **(g)** replication reconstituted in a test tube (This experiment was performed two times). For gel source data, see Supplementary Fig. 1.

Rad53 halts origin firing by targeting Sld3, as well as the Dbf4 subunit of DDK. In the Supplementary Results section, we show that Rad53 phosphorylates a segment of Sld3, involved in binding the A4 site, hence blocking origin firing by preventing Sld3 recruitment (**Supplementary Figs.4, 5**). This indicates that modulation of MCM engagement through phosphorylation by two different kinases has evolved to target three different interaction partners of the same Mcm4 binding pocket (Mcm4 itself, Dbf4^16,17,38^, and Sld3), which subsequently bind MCM on the path to origin activation)^9-13,33^.

### Sld3 tethers Cdc45 to MCM aided by Sld7

Opposite the A4 element, on the juxtaposed MCM hexamer (in *trans*), Cdc45 is contacted by a central alpha helical domain of Sld3 (amino acids 156-421), known as Cdc45-binding domain, or CBD^23^ (**Figs. 3a,b**). A first tethering element of Sld3 is provided by an alpha helix (residues 340-359), wedged in a hydrophobic groove within Cdc45. Mutational analysis previously showed that five of these residues (D344, D348, I352, I355 and L356) are essential for Cdc45 binding and cell viability (**Fig. 3c**). Cryo-EM density for the second tethering element was not as well resolved. To complement our atomic model, we docked the recently determined crystal structure of a yeast heterodimeric complex containing Cdc45 and Sld3 CBD, which provided an excellent fit into the pre-IC map^39^. According to the crystal structure, an alpha helix occupies the poorly defined density at the Sld3-Cdc45 interface, with Lys303 and Arg305 of Sld3 engaging in polar interactions with Glu239 and Ser647 of Cdc45 respectively (**Fig. 3d**). A 6x reverse charge mutation variant of Sld3, including Lys303Glu and Arg305Glu, impairs Cdc45 interaction and replication reconstituted in a test tube^23^. We conclude from these data that Cdc45 incorporation into the pre-IC involves the same interactions described for the isolated Sld3 CBD-Cdc45 heterodimer^39^.

**Figure 3.**
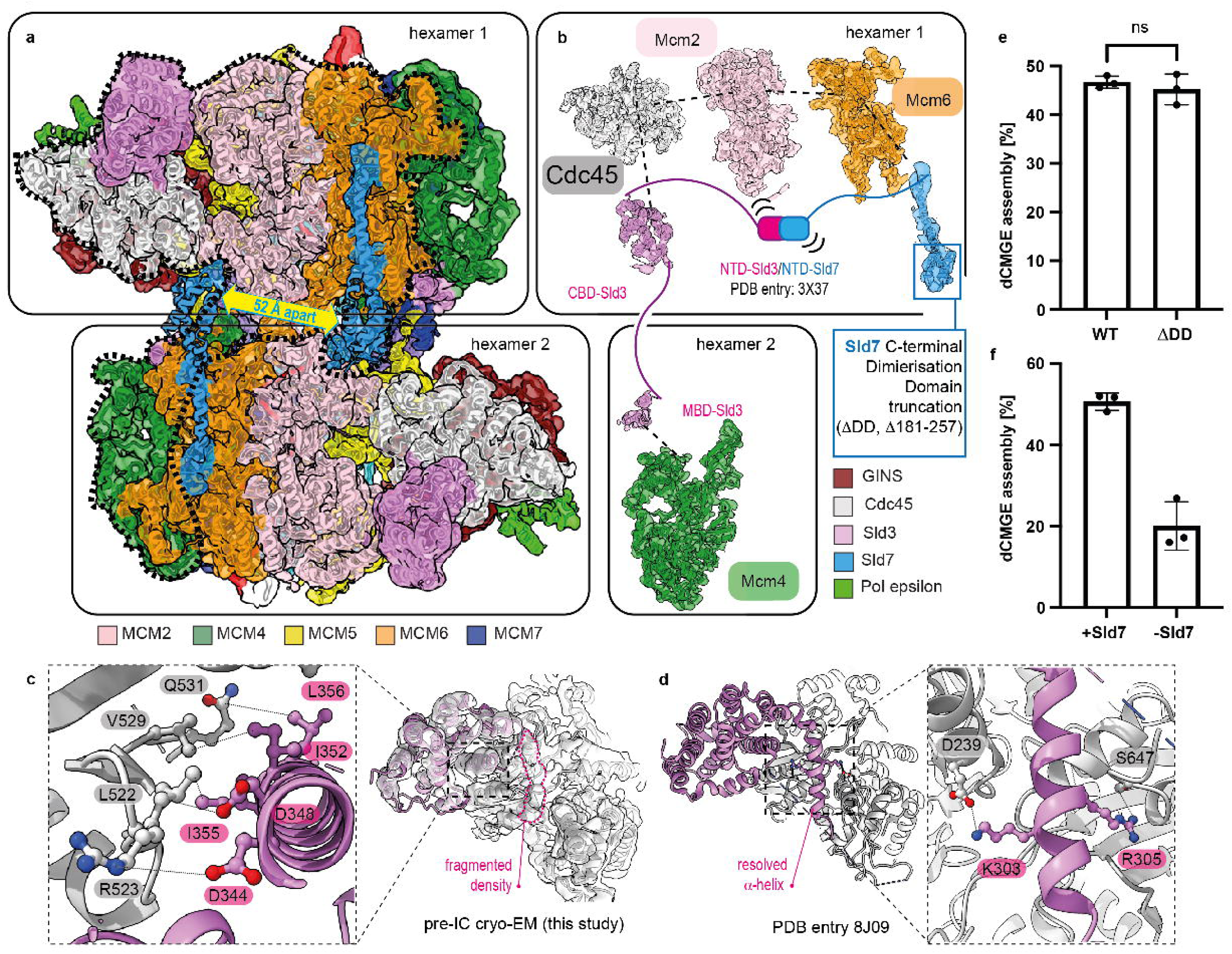
Sld3/7 recruit Cdc45 to MCM in the pre-IC. **(a)** Sld3 engages Mcm4 on one MCM hexamer and deposits Cdc45 onto the opposed MCM hexamer (marked by black dotted line). Two separate Sld7 molecules, distanced 52 Å apart, straddle across two Mcm6 subunits. **(b)** Interaction map of Sld3-Cdc45-Mcm2-Mcm6-Sld7 on one MCM hexamer and Sld3-Mcm4 on the opposing hexamer. Also refer to **Extended Data Figs. 3f-g**. While Sld3 is adjacent to Mcm4 on the opposed MCM ring, making C-terminal Sld3-Mcm4 engagement most likely, we cannot rule out that Sld3 could take a longer route wrapping around Mcm2,6 and Sld7 to connect Cdc45-binding and MCM-binding domains within the same MCM hexamer. **(c)** Cryo-EM density for Cdc45-Sld3. Detail of a first tethering point provided by an alpha helix element (residues 340-359). Density for a second tethering point is fragmented (dotted pink line). **(d)** Atomic model of the Cdc45-Sld3 determined by X-ray crystallography (8J09). Inset describes Sld3 residues whose mutations impair Cdc45 recruitment and replication. **(e)** Truncation of the Sld7 dimerisation domain (DD) does not impair dCMGE assembly (P-value determined by two-tailed t-test; *P*=0.4941; Error bars, mean ± s.d. This experiment was performed three times). **(f)** Omitting Sld7 allows dCMGE assembly, although at reduced efficiency (P-value determined by two-tailed t-test; *\*\*P*=0.0011; Error bars, mean ± s.d. This experiment was performed three times).

Sld3 was expressed and purified in complex with Sld7, a non-essential factor known to increase replication initiation efficiency^40,41^. We used AF3^37^ predictions to help locate cryo-EM density for Sld7 (amino acids 151-257). Sld7 features a 48-residues-long alpha helix, straddling the outer perimeter of the Mcm6 N-terminal and ATPase domains, and ending with a C-terminal helical bundle pointing towards the N-terminal domain of MCM where two hexamers meet. The local resolution of the pre-IC map was sufficient to allow model building of these Sld7 features. (**Figs. 3a,b**). Crystallographic evidence was previously used to propose that Sld7 might form a functional homodimer in solution, mediated by its C-terminal dimerisation domain (DD). This model supports a symmetric mechanism for the concomitant recruitment of two Cdc45 factors. While all pre-IC particles always contain two copies of Cdc45 (as well as GINS), we find that the two Sld7 DD domains do not engage in homo-dimerisation, rather they map 52 Å apart (**Fig. 3a**). To test the functional importance of the C-terminal module we generated an Sld3/7 variant bearing an Sld7 truncation of residues 181-257 (Sld7ΔDD, **Fig. 3b**). We performed nsEM analysis to show that Sld7ΔDD supports efficient dCMGE assembly (**Fig. 3e**, **Extended Data Figs. 3a-c**). Conversely, when Sld7 was dropped out, dCMGE formation occurred inefficiently (**Fig. 3f**, **Extended Data Figs. 3a,d,e**). Although we were unable to unambiguously visualise a direct interaction between Sld3 and Sld7 in our pre-IC cryo-EM map, a co-crystal structure of the N-terminal heterodimerisation domains exists^27^. AF3^37^ co-folding of Mcm2, Mcm6, Cdc45, Sld3 and Sld7 predicts this heterodimerisation complex to be tethered to the rest of the pre-IC via an unstructured element, connecting Sld3 CBD to the Mcm6 interacting helix of Sld7 (**Extended Data Fig. 3f**). This would explain why the Sld3/7 heterodimerisation element is not visible in our cryo-EM map.

### Dpb11 recruits GINS onto splayed DHs

Wedged in between the two splayed MCM rings we identified two copies of Dpb11 (BRCA1 C-terminus, BRCT1 and BRCT2 domains). Although they are proximal, these two Dpb11 subunits do not interact. Rather, they symmetrically bridge between two Mcm3 and Mcm7 subunits aligned across the two MCM rings of pre-IC (**Fig. 4a**). Despite the limited resolution of the interaction interface, we could recognise alpha helix 1 of BRCT1 directly engaging the Mcm7 NTI (**Fig. 4b**). AF3^37^ recapitulated the same interaction interface, allowing us to identify three Dpb11 candidate residues engaged in a network of polar and hydrophobic contacts with Mcm7 (**Fig. 4c**). To disrupt these contacts, we introduced three mutations in Dpb11 (K21E, K25E and I28E, hereafter Dpb11 3E) (**Extended Data Fig. 3h**). When using this Dpb11 variant, we observed a four-fold decrease in DH to pre-IC conversion efficiency, and a small drop in replication signal from reconstituted reactions (**Fig. 4d** and **Extended Data Figs. 3h-j**). Dpb11 sequesters Mcm7 NTI, which no longer protects the Mcm5 A domain of the opposed MCM hexamer in the DH^20^. This structural change exposes an epitope on the Mcm5 A domain, allowing GINS engagement via the Psf2 subunit^22^ (**Fig. 4e**). Dpb11 also directly engages GINS subunits Psf1, Psf2 and Sld5, via an alpha helical element (residues 273-295, **Fig. 4f**). Despite the poor sequence conservation, the same GINS-interaction (GINI) element had previously been identified in a comparative biochemical study of yeast Dpb11 and the orthologous TopBP1 protein of *Xenopus laevis*^42^. Indeed, a cryo-EM structure of a frog TopBP1-GINS complex revealed a structurally conserved GINI alpha helix serving the same function observed in our yeast pre-IC^43^. We introduced three mutations in Dpb11 GINI residues (I287S, W288A and K290E, Dpb11 3X) involved in GINS engagement (**Extended Data Fig. 3h**). Using this mutant protein in reconstitution experiments yielded a 4-fold drop in DH to pre-IC formation efficiency and a severe replication defect (**Figs. 4g,h** and **Extended Data Fig. 3k**). When combining the Mcm7 NTI interaction and GINI mutants (Dpb11 3E/3X) we observed an additive detrimental effect, with a replication product barely above background level (**Fig. 4h**). Our structure uncovers an unexpected function of Dpb11 acting across the pre-IC, through engagement of Mcm7 on one MCM hexamer and recruitment of GINS onto the opposed hexamer. Thus, just like DDK^14^ and Sld3 (described above), each copy of Dpb11 engages two opposed MCM hexamers that contain the symmetry to support bidirectional replication. Furthermore, in the Supplementary Results section, we explain how Dpb11 exerts its reported phospho-reader function. The BRCT1-2 epitope on Dpb11 is solvent exposed and poised to bind CDK-phosphorylated Sld3 (**Supplementary Figs. 6a-d**), providing another example of an essential, kinase-driven mechanism that triggers origin firing^10,11^.

**Figure 4.**
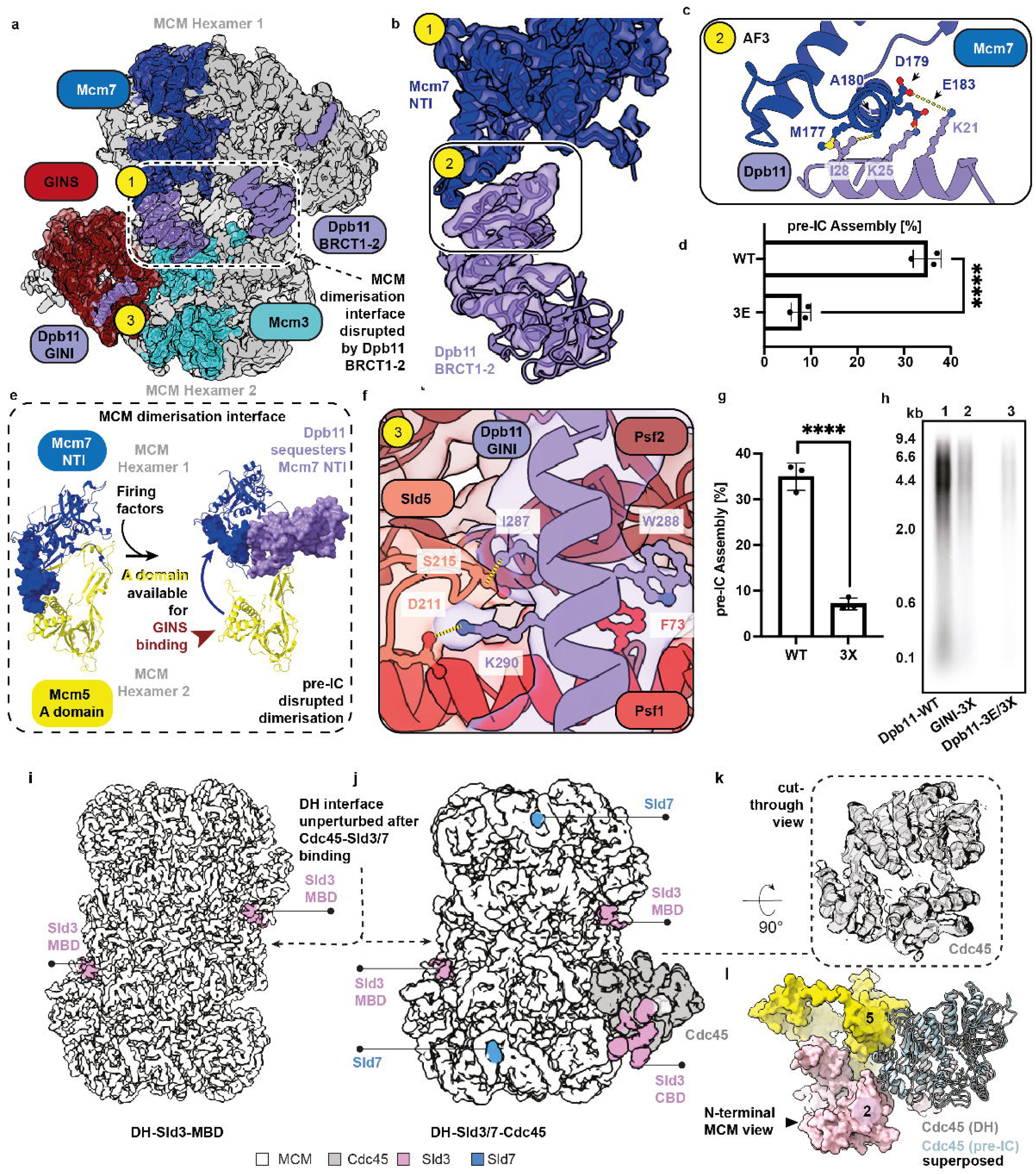
Structural role of Dpb11 in the pre-IC. **(a)** Dpb11 wedges itself between Mcm3 and Mcm7, at the interface between two MCM hexamers (yellow label 1). Dpb11 also engages GINS (yellow label 3). **(b)** Cryo-EM density with refined atomic model of Mcm7-Dpb11. Label 2 indicates Dpb11-Mcm7 interaction interface **(c)** AlphaFold3 prediction of Mcm7-Dpb11 interaction interface. **(d)** Mutating 3 Mcm7-interacting residues in Dpb11 causes a big drop in pre-IC complex formation, as observed by nsEM (P-value determined by one-way ANOVA; *\*\*\*\*P<*0.0001; Error bars, mean ± s.d. This experiment was performed three times). **(e)** Mcm5-Mcm7 compared in the DH and pre-IC structures. In the DH, the Mcm7 NTI protects the Mcm5 A domain. In the pre-IC, Mcm7 NTI is sequestered by Dpb11, rendering the Mcm5 A domain available for GINS binding. **(f)** Detailed view of the interaction between GINS and the GINI element of Dpb11. **(g)** A three-amino acid change in GINI impairs pre-IC assembly **(h)** and replication. (In (g), P-value determined by one-way ANOVA; *\*\*\*\*P*<0.0001; Error bars, mean ± s.d. This experiment was performed three times). Panels d and g show the same wild type condition because Dpb11 variants were probed within the same three replicate experiments. A mutant that also targets the Dpb11-Mcm7 interaction exacerbates the replication defect (This experiment was performed two times). For gel source data, see Supplementary Fig. 1. **(i)** Surface rendering of a DNA-loaded MCM double hexamer (DH) bound by the Sld3 MCM binding domain (MBD). **(j)** Surface rendering of the DH structure bound by Sld3-Sld7-Cdc45. Two copied of Sld3-7 and one copy of Cdc45 are visible. The Sld3 Cdc45 binding domain (CBD) recruits Cdc45 to MCM. **(k)** Detail of the cryo-EM density for Cdc45. **(l)** Cdc45 is slotted between the A domains of Mcm2 and Mcm5, in a position virtually identical to that observed in the pre-IC structure.

### What disrupts the DH interface

From the pre-IC structure alone, it is unclear whether Dpb11 binding breaks the DH interface or accesses a pre-formed gap, and if Mcm7-Sld3 engagement causes or is enabled by DH cracking. Previous reports indicate pre-IC formation involves two biochemically steps that can be staged^9,23,24,40^. To reconstitute the Cdc45-assembly step, we repeated the pre-IC formation reaction dropping out Sld2, Dpb11, Pol epsilon and GINS^30^ (**Extended Data Figs. 4a-d**), and analysed the reaction by cryo-EM. We solved a 3.1 Å resolution structure of a DH featuring Sld3 density at the A4 site, which possibly represents a first recruitment intermediate. In this structure Mcm7 NTI still tightly interacts with Mcm5, sealing the interface between the two hexamers (**Fig. 4i**). Following symmetry expansion, 3D classification and local refinement, we also solved a 3.7 Å resolution structure of DH-Sld3/7-Cdc45 (DH-3745, **Fig. 4j**). Sld3 CBD became visible, yet at lower local resolution, reflecting high flexibility compared to the pre-IC structure (**Extended Data Figs.4e-h,5** and **Extended Data Table 1**). Cdc45 slots in between the A domains of Mcm2 and Mcm5, occupying the same position observed in pre-IC and CMG (**Figs. 4k,l**). Sld7 was also observed to engage Mcm6, at the same ATPase site observed in the pre-IC (**Fig. 4j** and **Extended Data Fig. 3g**). Parallel 3D classification efforts provided some evidence for the flexible N-terminal domains connecting Sld3 and Sld7, compatible with previous X-ray work^27^ and the AF3^37^ prediction described above (**Extended Data Figs. 3f,g**). MCM in the DH-3745 structure is similar to a DDK-treated DH solved in ATP-containing buffer^14^, although nucleotide occupancy is different (**Supplementary Figs. 7a,b**)^38^. The MCM dimerisation interface of the DH does not change upon Sld3/7-Cdc45 binding, and an overlay of DH with Mcm7-Dpb11 extracted from the pre-IC structure reveals extensive steric clashes with Mcm5 (**Supplementary Fig. 7c**). We conclude that the separation between the two MCM rings must involve Sld2, Dpb11, GINS, Pol epsilon recruitment or a subset of these factors.

Our EM imaging of reconstituted Cdc45 recruitment by Sld3/7 reveals one or two Cdc45 molecules engaged to DHs at any one time (**Extended Data Figs. 4b,d**), suggesting that Cdc45 is recruited to each MCM hexamer in independent events. This is compatible with single-molecule measurements using yeast proteins^25^ and different from single-molecule observations with frog egg extracts. In fact, in the Xenopus system, Cdc45 is recruited to DHs synchronously, via a mechanism that is yet to be uncovered^44^.

In the Supplementary Results, we discuss the changes in DNA engagement triggered upon transition between DH-3745 and pre-IC (**Supplementary Figs. 8a,b**).

### How CMG assembly factors are ejected

The only Pol epsilon element visible in the pre-IC is the N-terminal domain of Dpb2 (**Extended Data Fig. 6a**). This observation caught our attention for two reasons. First, the isolated N-terminal Dpb2 was observed to support CMG assembly upon depletion of endogenous Dpb2 in yeast cells^18^. Second, according to structural studies on CMGE bound to an artificial DNA fork, N-terminal Dpb2 is the only Pol epsilon element detected in CMG when the Mcm2-5 site is nucleotide-free (just like in pre-IC, **Supplementary Fig. 7b, Extended Data Fig. 6b**)^45^. Instead, when Mcm2-5 is ATP-bound, the non-catalytic portion of Pol epsilon can be averaged in full, stably anchored to MCM^30,45,46^ (as seen in ATP-dCMGE, **Extended Data Figs. 6c,d**). Inspired by these observations, we hypothesised that not only Dpb2, but the full Pol epsilon complex might be tethered to CMG in our pre-IC structure, and it would become fully MCM-engaged (visible) when ATP is added. To test our hypothesis, we assembled the pre-IC in the absence of nucleotide, as described above, and purified this complex washing away excess Pol epsilon (**Extended Data Fig. 6e**). After elution and incubation with or without ATP we imaged particles by nsEM. Only when ATP was supplemented, did we recognise the non-catalytic Pol epsilon decoration characteristic of ATP-dCMGE complexes. As observed before, the two CMGE particles were present in both in the cis and trans configuration (**Extended Data Fig. 6f**)^30^.

ATP-dependent pre-IC to dCMGE transition suggests a mechanism for how ATP binding at an origin of replication might stabilise topological binding of CMG to DNA, making it resistant to high-salt washes^24^. In fact, while GINS and Cdc45 latch across the Mcm2-5 gate on the N-terminal MCM side of CMG, fully engaged Pol epsilon stabilises the C-terminal MCM side, by securing GINS and Cdc45 to the Mcm2-5-3 ATPase domains^46^. An ancillary role of Pol epsilon in CMG assembly is discussed in **Supplementary Figs. 7d,e**.

Tight Pol epsilon engagement to C-terminal CMG in the ATP-dCMGE complex also addresses a key unknown in replication origin activation: how assembly factor eviction is triggered through ATP binding by MCM^24,30^. In fact, Cdc45 engagement by C-terminal Pol2 sterically clashes with the Sld3 CBD, providing a mechanism for Sld3 ejection (**Extended Data Fig. 6g**). The ability of two ATP-CMGEs to rotate with respect to one another^30^ provides another means for the release of CMG assembly factors. Indeed, as soon as two CMGs visit a cis dCMGE configuration, Dpb11 would be dislodged from its position, nestled between Mcm3 and Mcm7 across the two hexamers, (**Fig. 4a**). CMG rotation would also lead to eviction of Sld3 itself, as Sld3 binds Cdc45 on one CMG and Mcm4 in the opposed CMG across the pre-IC (**Fig. 3b**).

### Sld2 is essential after CMGE assembly

In the pre-IC cryo-EM map, density for all factors previously implicated in CMG assembly can be identified, except for one, Sld2. This was not surprising, given that phospho-Sld2 is known to engage a flexibly tethered region of Dpb11^34^, which is not resolved in our pre-IC structure. To establish whether Sld2 is indeed required for GINS recruitment, as commonly accepted, we assembled pre-IC as described above and analysed the reaction by nsEM, either using the full complement of firing factors or dropping out individual components. While ppSld3/7, GINS and Cdc45 were all required for pre-IC formation (**Supplementary Figs. 7d,e**), dropping out the Sld2 8D phosphomimetic variant still yielded recognisable pre-IC averages, although DH-to-pre-IC conversion rate dropped by roughly one third (from 28% to 20%, based on 16,318 averageable particles for Sld2 8D, 22,234 for the dropout, each over three biological replicates, **Figs. 5a-c**). Similarly reduced efficiency (based on 20,030 averageable particles) was obtained when swapping Sld2 8D for non-phosphorylated, wild type Sld2 (**Figs. 5a-c**). When repeating these experiments in the presence of ATP, dCMGE^30^ could be assembled with robust efficiency, reflecting the increased stability provided by nucleotide binding^24^ (analysis based on 55,833 averageable particles over the three conditions, **Figs. 5d,e**).

**Figure 5.**
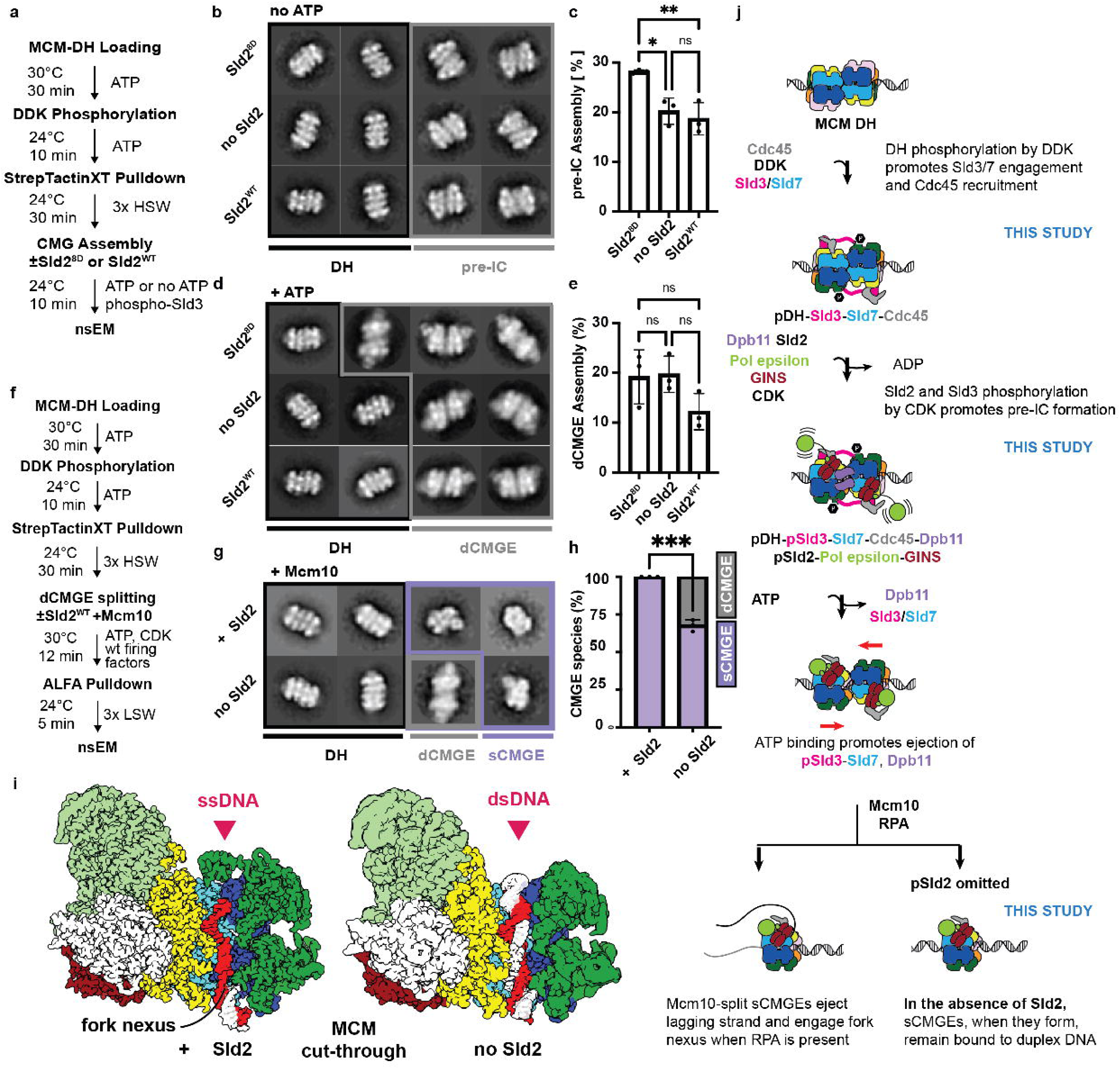
Exploring the role of Sld2 in CMG assembly. **(a)** Flow chart of pre-IC and dCMGE formation with or without Sld2, 8D variant or wt. **(b)** Representative 2D averages of pre-IC assembly reaction. **(c)** Bar graphs of pre-IC formation with or without Sld2, 8D variant or wild type. This experiment was performed three times. One-way ANOVA P Values: 8D vs noSld2 *P=0.0160; 8D vs wt Sld2 **P=0.0070; noSld2 vs Sld2-WT P=0.7293; Error bars, mean ± s.d. **(d)** Representative 2D averages of dCMGE assembly reaction. (**e**) Bar graphs for dCMGE formation with or without Sld2, 8D variant or wild type. This experiment was performed three times. One-way ANOVA P values: 8D vs noSld2 P=0.9875; 8D vs wt Sld2 P=0.1981; noSld2 vs Sld2-WT P=0.1635; Error bars, mean ± s.d. **(f)** Flow chart of dCMGE splitting by Mcm10 in the presence or absence of Sld2. **(g)** Representative 2D averages of dCMGE splitting reaction. **(h)** Bar graph showing that Sld2 is not strictly required to split dCMGE into single CMGEs, however its absence negatively affects splitting efficiency. This experiment was performed three times. Two-tailed t test, ***P=0.0001; Error bars, mean + s.d. **(i)** Cut-through side view of a single CMGE assembled in the presence of Mcm10 and RPA, with or without Sld2. In the presence of Sld2, sCMGE engages the fork nexus at the N-terminal front. Only the leading strand template enters the MCM pore. In the absence of Sld2, duplex DNA is observed running through the entire length of the MCM pore. The structure assembled with Sld2 is coloured according to a newly built atomic model. The structure assembled without Sld2 is coloured according to a previously published dsDNA-CMGE atomic model (PDB entry 7HQS). **(j)** Cartoon representation of origin activation, showing that Sld2 plays an essential origin activation function downstream of CMG formation.

So far, we have shown that Sld2 can stimulate but is not needed for pre-IC assembly or dCMGE formation. Previous biochemical reconstitution however established that phospho-Sld2 is strictly required for origin dependent DNA replication^40^. Given that we can reproduce the observation with the same purified proteins used in our nsEM experiments (*not shown*), we conclude that an essential function of Sld2 during origin activation must be performed downstream of dCMGE assembly. To address what step after CMGE formation requires phospho-Sld2, we reconstituted Mcm10-dependent dCMGE splitting from purified DDK-phosphorylated DHs, in the presence of CDK and RPA, with or without wild type Sld2. We further purified this preparation using paramagnetic beads to pull on an ALFA-tagged covalently linked methyltransferase roadblock capping the ends of the double helix, to ensure that all helicase particles imaged were engaged to DNA (**Fig. 5f**). According to our nsEM analysis, when Sld2 was included, only DHs and sCMGEs were observed (based on 28,577 averageable particles). This recapitulates previous observations that all dCMGEs assembled are split into sCMGE when Mcm10 is added^47^. When Sld2 was omitted, instead, we observed a mixture of dCMGEs (32%) and sCMGEs (68%, based on 25,386 averageable particles over three repeats, **Figs. 5g,h**). We then asked whether sCMGE assembled on origin DNA without Sld2 is different compared to sCMGE obtained with the complete set of firing factors. To address this, we analysed the two reactions by cryo-EM.

When Sld2 and RPA were present, the 3D structure revealed sCMGE engaged to the fork nexus, with single-stranded DNA running through the MCM central channel (no sCMGE engaged to duplex DNA could be identified in this condition, when further 3D classification was attempted). When Sld2 was dropped out, however, sCMGE was found to be duplex-DNA bound (no sCMGE engaged to single-stranded DNA could be identified). Engagement of the double helix in this context appears identical to what was observed in dCMGE^30^ (PDB entry 7HQS, **Fig. 5i**, **Extended Data Figs.7 and 8**, **Extended Data Table 2**), or sCMGE at termination^48^. Our data imply that Sld2 plays an essential role in lagging strand ejection after dCMGE formation.

## Discussion

The factors involved in eukaryotic DNA replication initiation are largely conserved from yeast to humans^49^. Exceptions exist, both in terms of function and identity of the proteins involved. For example, all six subunits of the Origin Recognition Complex, the loader of the MCM helicase, are essential for helicase loading in yeast, but not in humans^50,51^. Likewise, Mcm10 is required to split the dCMGE and establish bidirectional replication in yeast, but it is dispensable in *C. elegans*^52^. The picture becomes more complicated when one specific gene is lost during evolution, as other factors must be repurposed to fulfil the missing function. This is the case for DONSON, which is present in most eukaryotes but has been lost in yeast. Recent studies in metazoans established that DONSON performs an essential function during CMG assembly, by delivering GINS to MCM^52-55^. In yeast, the GINS delivery function has been assigned to Sld2, based on observations that GINS is not detected on chromatin when Sld2 is absent^10,26^.

Does DONSON indeed perform the same function as yeast Sld2, as recently proposed? Phylogenetic analysis quickly unveils a complex scenario, casting doubt on the matter. For example, *C. elegans* encodes both DONSON and yeast-like SLD2^56^, while most of the other DONSON-containing eukaryotes encode the RECQL4 helicase^57^, a multidomain protein featuring an Sld2-homology domain. This domain alone is essential for origin activation in *Xenopus laevis* but functions downstream of, not during, CMG assembly^58-61^. Thus, a more appropriate question is: has the role of Sld2 (metazoan RECQL4) been repurposed in yeast to make up for the loss of DONSON?

Our results do not fully support this scenario. In agreement with previous work, we find that Sld2 is involved in the efficient delivery of GINS to MCM, as observed in the pre-IC assembly step of CMGE biogenesis. However, Sld2 is not absolutely required for making a stable dCMGE complex. Instead, it plays an essential role after holohelicase assembly. In fact, we find that dCMGE splitting is defective without Sld2 and helicases that still mature to the sCMGE state, in these conditions, fail to transition to engaging single-stranded DNA. They remain duplex-DNA bound, instead. We conclude that the processes of double CMGE splitting, MCM gate opening and controlled lagging strand ejection must be tightly coupled and all require Sld2 to function productively (**Fig. 5j**). In this respect, we note that, according to Western blot analysis of the same reconstituted system the amount of CMG retained on DNA in the presence of Mcm10 is markedly reduced when Sld2 is omitted^40^. Does this imply that selective strand ejection fails without Sld2, such that not only the lagging but also the leading strand DNA can be evicted, putting CMGEs at risk of falling off the double helix? Is the loss of a sCMGE partner the reason why orphan sCMGEs remain bound to duplex and do not transition to single-stranded DNA? In other words, can lagging-strand ejection only occur when two CMGEs cross paths? From an evolutionary perspective, is the lagging strand ejection function of Sld2 conserved in RECQL4^58-61^? Further work is needed to address these topics.

Another question remains to be addressed. What yeast factor, if not Sld2, serves the role of DONSON during CMG formation? When inspecting the yeast pre-IC and the *X. laevis* double CMG-DONSON complex^62^ we noticed a tantalising similarity. Just like frog DONSON, yeast Dpb11 sits across the N-terminal homo-dimerisation domain of MCM, and it concomitantly engages GINS and Mcm3. Based on this observation, we suggest that Dpb11 might at once play the essential phospho-reader role of its known ortholog, TopBP1^63^, as well as the GINS-recruitment function of DONSON.

## Materials and Methods

### Protein expression and purification

MH, ORC, Cdc6, Mcm2-7/Cdt1, DDK, CDK, (yeast-expressed) Sld3/7, Cdc45, GINS, (yeast-expressed) Pol epsilon, Mcm10, Replication Protein A (RPA), Topoisomerase I (TopoI), Pol α and Rad53 were expressed and purified as previously described^24,30,40,46,64-67^. All mutant constructs were expressed and purified following the same protocol as the wild-type protein unless stated otherwise. All buffers described below are also reported in **Supplementary Table 1**.

### Cell lines

Sf21 insect cells were obtained in-house from the Cell Services Science and Technology Platform. These cells were not authenticated and tested negative for mycoplasma contamination.

### Cloning, expression and purification of Twin-Strep-tagged Sld3/7

Codon-optimized gene blocks (IDT) encoding *S. cerevisiae* Sld3 in frame with a Tobacco Etch Virus (TEV) protease cleavage site and a C-terminal Twin-Strep-tag (TST) as well as *S. cerevisiae* Sld7 were inserted into GoldenBac shuttle vectors pGB-01;02 and pGB-02;03. Subsequently, Sld3-TEV-TST and Sld7 expression cassettes were subcloned into pGB-dest using GoldenBac assembly^68^ and transformed into electro-competent EMBacY cells (Geneva Biotech). Cells were screened by blue-white selection for successful bacmid integration and selected colonies were grown overnight at 37°C. Cells were harvested by centrifugation and bacmids purified by isopropanol precipitation. 45 µg of bacmid DNA were mixed thoroughly with 13.5 µL FuGENE HD Transfection Reagent (Promega) and 450 µL Sf-900 III SFM media and incubated for 30 minutes at room temperature. 200 µL transfection mix were added dropwise to 2 mL Sf21 insect cells seeded at 0.5 million cells/mL in a 6-well plate. Plates were incubated for 3 – 5 days at 27°C in a wet-towel box. Efficient transfection was monitored via YFP fluorescence. Adherent cells were resuspended and supernatant containing P0 baculovirus was collected. To increase the multiplicity of infection, 46 mL of Sf21 insect cells at 0.5 million cells/mL were inoculated with 4 mL P0 baculovirus in suspension and cultured at 27°C shaking at 120 rpm. After viability had dropped below 90%, cells were pelleted by centrifugation at 380 x g for 15 minutes at 4°C, supernatant containing the amplified P1 virus was sterile-filtered (0.22 µm pore size) and stored at 4°C. 1L Sf21 insect cells were seeded at 1 million cells/mL in Sf-900 III SFM media and infected with 0.5 % v/v P1 virus. Cells were harvested 48 hours after baculovirus-induced cell-cycle arrest by centrifugation for 15 minutes at 180 x g at 4°C, pellets were flash-frozen in liquid nitrogen and stored at -80°C.

Cells were thawed and resuspended in 50 mL Buffer A (25 mM HEPES-KOH pH 7.5, 500 mM KCl, 10% v/v glycerol, 0.02% w/v NP-40, 1 mM EDTA, 1 mM DTT) supplemented with 1 cOmplete EDTA-free protease inhibitor tablet (Merck) and 0.7 mM Phenylmethylsulphonyl fluoride (PMSF), then lysed by sonication on ice for 2 minutes (1 second pulse-on, 4 seconds pulse-off). The lysate was clarified by ultracentrifugation for 1 hour at 45,000 rpm in a Ti45 rotor (Beckman) at 4°C, the supernatant was mixed with 2.4 mL Bio-Lock (IBA) reagent and applied onto 1 mL pre-equilibrated StrepTactinXT Superflow HighCapacity resin in a gravity column. The resin was washed with 100 mL Buffer A and 10 mL Buffer A supplemented with 2 mM ATP and 10 mM MgCl_2_. Protein was eluted with 10 mL Buffer A supplemented with Buffer BXT (IBA). The eluate was pooled, concentrated and loaded onto a Superdex 200 Increase 10/300 GL column (Cytiva) equilibrated in Buffer A. Gel-filtered Sld3/7 was concentrated to approximately 1.5 mg/mL, aliquoted and flash-frozen in liquid nitrogen.

### Cloning, expression and purification of Dpb11

Codon-optimized *S. cerevisiae* Dpb11 followed by a 3C protease cleavage site and a C-terminal 3xFLAG tag was subcloned into a pGB-04;05 shuttle vector and subsequently transformed into electro-competent EMBacY cells. Bacmid and baculoviruses were prepared as described above, 1L Sf21 insect cells at 1 million cells/mL were infected with 0.5% v/v P1 virus and harvested 48 hours after cell-cycle arrest.

The cell pellet was resuspended in 50 mL Buffer A supplemented with 1 cOmplete EDTA-free protease inhibitor tablet (Merck)+ 0.7 mM PMSF, lysed by sonication on ice for 2 minutes (1 second pulse-on, 4 seconds pulse-off) and ultracentrifuged at 45,000 rpm at 4°C for 1 hour. The soluble phase was mixed with 2.4 mL Bio-Lock Reagent and passed through 1 mL pre-equilibrated Anti-FLAG M2 Affinity Gel (Sigma) in a gravity column. The column was washed with 150 mL Buffer A and 10 mL Buffer A supplemented with 2 mM ATP and 10 mM MgCl_2_. To elute bead-bound protein, the beads were resuspended in 5 mL Buffer A supplemented with 0.5 mg/mL 3xFLAG peptide and incubated for 5 minutes, after which the flow-through was collected. The eluate was diluted to 150 mM KCl and loaded onto a 1 mL Mono S 5/50 column (Cytiva) equilibrated in Buffer B (25 mM HEPES-KOH pH 7.5, 150 mM KCl, 10% v/v glycerol, 0.02% w/v NP-40, 1 mM EDTA, 1 mM DTT). After washing the column with 10 mL Buffer B, Dpb11 was eluted with a linear gradient from 150 – 1000 mM KCl in Buffer B over 20 column volumes. Fractions containing pure Dpb11 were pooled and dialysed against Buffer C (25 mM HEPES-KOH pH 7.5, 300 mM KOAc, 10% v/v glycerol, 0.02% NP-40, 1 mM EDTA, 1 mM DTT) at 4°C overnight while stirring. Subsequently, Dpb11 was concentrated to approximately 0.5 mg/mL, aliquoted and flash-frozen in liquid nitrogen.

Dpb11 mutants containing charge-reversal substitutions (Dpb11^3E^, Dpb11^3X^, Dpb11^3E/3X^) were purified using gel filtration instead of cation exchange chromatography. The reason for this alteration was the predicted isoelectric point of the mutant proteins, which matches the pH of Buffers B and C. After FLAG affinity purification, the eluate was concentrated and loaded onto a Superdex 200 Increase 10/300 GL column equilibrated in Buffer C. Dpb11-containing fractions were pooled, concentrated to approximately 0.5 mg/mL, aliquoted, and flash-frozen in liquid nitrogen.

### Cloning, expression and purification of Sld2

An expression cassette encoding *S. cerevisiae* Sld2 in frame with an N-terminal VNp6 peptide tag^69^ and a retro-protein XXA solubility tag^70^ as well as a C-terminal Twin-Strep-tag was subcloned into a pET303 backbone and transformed into bacterial T7 Express cells. Multiple colonies were picked to inoculate 4x 1L LB + 100 µg/mL Carbenicillin and incubated static overnight at 37°C. The next morning, cultures were transferred to 30°C and grown to an OD_600_ of 0.8 shaking at 200 rpm. 80 µM IPTG were added to each flask to induce Sld2 expression for 21 hours at 30°C at 200 rpm. Subsequently, cells were pelleted by centrifugation at 4,000 x g for 10 minutes at 4°C, flash-frozen and stored at -80°C.

Cells were thawed and resuspended in 100 mL Buffer D (25 mM HEPES-KOH pH 7.5, 800 mM KCl, 10% v/v glycerol, 1M Sorbitol, 2 mM ATP, 10 mM MgCl_2_, 0.02% v/v NP-40, 0.1% w/v Tween-20, 1 mM DTT) + 2 cOmplete EDTA-free protease inhibitor tables (Merck) and 0.7 mM PMSF, then sonicated on ice for 2 minutes (5 second pulse-on, 5 seconds pulse-off). The lysate was clarified by centrifugation in a JA-25.50 rotor at 18,000 rpm for 20 minutes at 4°C, the supernatant was applied onto a gravity column packed with 1 mL StrepTactinXT Superflow HighCapacity resin equilibrated in Buffer D. The resin was washed with 75 mL Buffer D followed by 25 mL Buffer E (25 mM HEPES-KOH pH 7.5, 500 mM NaCl, 10% v/v glycerol, 0.02% w/v NP-40, 1 mM EDTA, 1 mM DTT), after which Sld2 was eluted by passing 10 x 1 mL Buffer E supplemented with Buffer BXT through the resin. The highest-concentrated fraction was identified by SDS-PAGE (Coomassie staining) and dialysed in Buffer F (25 mM HEPES-KOH pH 7.5, 700 mM KOAc, 40% v/v glycerol, 0.02% w/v NP-40, 1 mM EDTA, 1 mM DTT) for 4 hours at 4°C. Sld2 was aliquoted at approximately 0.8 mg/mL and flash-frozen in liquid nitrogen. For the phosphomimetic Sld2 8D variant, aspartate substitutions were introduced at the following residues: Threonine 84, Serine 100, Serine 128, Serine 138, Threonine 168, Serine 172, Serine 188, and Serine 208.

### Cloning, expression and purification of ALFA-tagged Pol epsilon

Codon-optimized *S. cerevisiae* Pol2-3xFLAG, Dpb2, Dpb3 and Dpb4-ALFA were subcloned into GoldenBac shuttle vectors and assembled into a co-expression plasmid, pGB-dest-PolE as described for Sld3/7 above. Similarly, electro-competent EMBacY cells were transformed with pGB-dest-PolE to prepare bacmids and generate a P1 baculovirus as described above. 1 billion Sf21 insect cells were seeded in 1L Sf900-III SFM media and infected with 0.5% v/v P1 baculovirus, incubated at 27°C at 120 rpm, and harvested 48 hours after cell-cycle arrest by centrifugation at 180 x g at 4°C for 15 minutes. The cell pellets were flash-frozen in liquid nitrogen and stored at -80°C. To purify ALFA-tagged Pol epsilon, the cell pellets were resuspended in 50 mL Buffer G (25 mM HEPES-KOH pH 7.6, 400 mM KOAc, 10% v/v glycerol, 2 mM DTT) supplemented with 1 cOmplete EDTA-free protease inhibitor tablet (Merck) and lysed by sonication on ice for 2 minutes (1 second pulse-on, 4 seconds pulse-off). The lysate was clarified by ultracentrifugation at 45,000 rpm for 45 minutes at 4°C in a Ti45 rotor, the supernatant was passed twice through a column packed with 1 mL Anti-FLAG M2 affinity gel equilibrated in Buffer G. The column was washed with 150 mL Buffer G and 20 mL Buffer G + 2 mM ATP and 10 mM Mg(OAc)_2_. Protein was eluted by incubating the resin three times in 5 mL Buffer G + 0.5 mg/mL 3xFLAG peptide for 5 minutes and collecting the flow-through. The eluate was loaded onto a 5 mL Heparin HP column (Cytiva) and eluted over 25 column volumes with a linear gradient from 400 – 1000 mM KOAc in Buffer G. Fractions containing Pol epsilon were concentrated and gel filtered onto a HiLoad 16/60 Superdex 200 pg column (Cytiva). Pol epsilon was concentrated to approximately 1 mg/mL, aliquoted and flash-frozen in liquid nitrogen.

### Cloning, expression and purification of Sic1

T7 Express cells (NEB) were transformed with hexahistidine-tagged *S. cerevisiae* Sic1*^71^.* Transformant colonies were incubated overnight in 100 mL LB supplemented with 100 µg/mL Carbenicillin at 37°C shaking at 200 rpm. 2L LB media + 100 µg/mL Carbenicillin were inoculated with 1% v/v dense overnight culture and grown to an optical density at 600 nm (OD_600_) of 0.6 at 37°C and 200 rpm. Expression of Sic1 was induced by adding 0.5 mM Isopropyl β-D-1thiogalactopyranoside (IPTG), after which the cultures were incubated for 3 hours at 37°C at 200 rpm. Cells were harvested by centrifugation at 4,000 x g for 10 minutes at 4°C, the cell pellet was flash-frozen in liquid nitrogen and stored at -80°C.

The pellet was dissolved in 100 mL Buffer H (25 mM HEPES-KOH pH 7.5, 500 mM NaCl, 10% v/v glycerol, 1 mM EGTA, 0.2% w/v Triton X-100, 0.5 mM TCEP, 10 mM Imidazole) + 2 cOmplete EDTA-free protease inhibitor tablets (Merck) and 0.7 mM PMSF in a beaker. 0.2 mg/mL Lysozyme were added and incubated with the cell suspension for 10 minutes while stirring. Cells were then sonicated on ice for 5 minutes (2 seconds pulse-on, 5 seconds pulse-off), the debris was removed by centrifugation at 20,000 rpm in a JA25.50 rotor for 30 minutes at 4°C. 2 mL Ni-NTA beads (Quiagen) were equilibrated in Buffer H and rotated with the clarified lysate for 1 hour at 4°C. Subsequently, beads were collected in a gravity column and washed with 150 mL Buffer H and 20 mL Buffer H supplemented with 2 mM ATP and 10 mM MgCl_2_. Protein was eluted with 15 mL Buffer H supplemented with 250 mM Imidazole, concentrated and gel-filtered on a HiLoad 16/60 Superdex 75 pg column (Cytiva) equilibrated in Buffer I (25 mM HEPES-KOH pH 7.5, 5% v/v glycerol, 5 mM MgCl_2_, 0.5 mM EDTA, 0.5 mM TCEP) supplemented with 200 mM NaCl. Gel filtration did not yield pure Sic1. Consequently, peak fractions were loaded onto a 1 mL Mono S column (Cytiva) and washed with 10 column volumes Buffer I supplemented with 50 mM NaCl. Sic1 was eluted with a linear gradient of 50 – 1000 mM NaCl in Buffer I over 30 column volumes, fractions containing pure Sic1 were dialysed at 4°C in Buffer I + 200 mM NaCl for 3 hours under agitation. Sic1 was concentrated to 10.8 mg/mL and flash-frozen in liquid nitrogen.

### Cloning, expression and purification of Twin-Strep-tagged Sumo-Mcm10

*S. cerevisiae* Mcm10 was cloned in frame with an N-terminal 10xHis-SUMO cassette and a C-terminal Twin-Strep-tag and transformed into Rosetta 2 pLysS cells. Multiple colonies were picked and grown overnight at 37°C in 100 mL LB supplemented with 100 µg/mL Carbenicillin and 33 µg/mL Chloramphenicol. 6x 1L LB supplemented with 100 µg/mL Carbenicillin and 33 µg/mL Chloramphenicol were each inoculated with 10 mL dense overnight culture and grown to an OD_600_ of 0.7 at 37°C and 200 rpm. Over-expression was induced by addition of 0.5 mM IPTG and continued for 16 hours at 16°C. Cells were collected by centrifugation at 4,000 x g for 10 minutes at 4°C, flash-frozen in liquid nitrogen and stored at -80°C. The cell pellet was resuspended in 230 mL Buffer J (25 mM HEPES-KOH pH 7.6, 500 mM NaCl, 10% v/v glycerol, 1 mM EDTA, 0.05% w/v Tween-20, 1 mM DTT) supplemented with 4 cOmplete EDTA-free protease inhibitor tablets (Merck), then lysed by sonication on ice for 5 minutes (2 seconds pulse-on, 5 seconds pulse-off). The lysate was clarified by centrifugation in a JA25.50 rotor at 20,000 rpm for 30 minutes at 4°C, the supernatant was loaded onto a 1 mL cOmplete His-Tag Purification column (Merck) installed in tandem with a 1 mL StrepTactinXT 4Flow high capacity column (IBA), both pre-equilibrated in Buffer J. The columns were washed with Buffer J until the absorbance at 280 nm returned to its base line signal, after which protein was eluted from the His-Tag Purification column into the connected StrepTactinXT column with 9 mL Buffer J supplemented with 200 mM Imidazole. The His-Tag Purification column was disconnected, the StrepTactinXT column was washed with 9 mL Buffer B, and finally eluted with 9 mL Buffer K (25 mM HEPES pH 7.6, 300 mM NaCl, 10% v/v glycerol, 0.05% w/v Tween-20, 1 mM DTT, 5 mM Desthiobiotin). Mcm10-containing fractions were pooled and dialysed overnight into Buffer L (25 mM HEPES pH 7.6, 200 mM NaCl, 20% v/v glycerol, 0.05% w/v Tween-20, 1 mM EDTA, 2 mM DTT) at 4°C while stirring. Mcm10 was aliquoted at a concentration of approximately 0.5 mg/mL and flash-frozen in liquid nitrogen.

### Preparation of MH-conjugated ARS1 DNA template

A 168-bp DNA template containing the *S. cerevisiae* origin of replication ARS1, flanked by two MH recognition sites was generated by PCR and purified as previously described^72,73^. The DNA template was covalently tethered to either Twin-strep-tagged MH or tandem ALFA/Twin-Strep-tagged MH using previously established protocols^30^ (Pühringer et al., *in preparation*).

#### S-CDK Prephosphorylation of Sld3/7

Sld3/7 prephosphorylation by S-CDK was modified from previous protocols^24^. In short, 900 nM Twin-Strep-tagged Sld3/7 were phosphorylated by 100 nM S-CDK in Buffer M (40 mM HEPES-KOH pH 7.5, 310 mM K-Glutamate, 10 mM Mg(OAc)_2_, 10% v/v glycerol, 0.02% w/v NP-40, 1 mM DTT, 2 mM ATP, 0.4 mg/mL BSA) in a total volume of 100 µL for 8 minutes at 24°C and 1,250 rpm. Phosphorylation was stopped by adding 2.2 µM Sic1 to the reaction and incubating it for 2 minutes at 24°C and 1,250 rpm. 500 mM KCl was added to the reactions, which were then bound to 10 µL MagStrep “type3” XT slurry (IBA) equilibrated in Buffer N (25 mM HEPES-KOH pH 7.5, 500 mM KCl, 5 mM Mg(OAc)_2_, 10% v/v glycerol, 0.02% w/v NP-40, 1 mM DTT). After 30 minutes of incubation at 24°C and 1,250 rpm, beads were washed 5x with 200 µL Buffer K, prephosphorylated Sld3/7 (ppSld3/7) was eluted in 10 µL Buffer N + 25 mM D-Biotin for 10 minutes at 24°C and 1,250 rpm, aliquoted, flash-frozen in liquid nitrogen and stored at -80°C. Successful phosphorylation and approximate yield were estimated by SDS-PAGE. For cryo-EM experiments, ppSld3/7 was prepared in parallel and not flash-frozen prior to its use.

### Rad53 Prephosphorylation of Sld3/7

Rad53-prephosphorylated Sld3/7 was prepared by incubating 700 nM Twin-Strep-tagged Sld3/7 with 356 nM Rad53 in 100 µL Buffer M for 30 minutes at 30°C and 1,250 rpm. Subsequently, Rad53-prephosphorylated Sld3/7 was purified via StrepTactinXT affinity purification exactly as described for CDK-prephosphorylated Sld3/7 above.

### In vitro ATP-dCMGE assembly

ATP-bound dCMGE complexes were assembled as previously described^30,47^ with minor modifications. First, MCM-DHs were loaded for 30 minutes at 30°C and 1,250 rpm by co-incubating 20 nM MH-conjugated ARS1 with 52 nM ORC, 52 nM Cdc6, and 110 nM Mcm2-7/Cdt1 in 100 µL Buffer O (25 mM HEPES-KOH pH 7.5, 100 mM K-Glutamate, 10 mM Mg(OAc)_2_, 1 mM ATP, 0.02% w/v NP-40). Afterwards, loaded MCM-DHs were phosphorylated with 80 nM DDK at 24 °C and 1,250 rpm for 10 minutes, then bound for 30 minutes at 24°C and 1,250 rpm to 5 µL MagStrep “type3” XT slurry equilibrated in Buffer P (25 mM HEPES-KOH pH 7.5, 100 mM K-Glutamate, 10 mM Mg(OAc)_2_, 0.02% w/v NP-40). The beads were washed 3x with 200 µL Buffer Q (25 mM HEPES-KOH pH 7.5, 500 mM NaCl, 5 mM Mg(OAc)_2_, 0.02% w/v NP-40) and 1x with 200 µL Buffer P, then eluted in 20 µL Buffer O + 25 mM D-Biotin for 15 minutes at 24 °C and 1,250 rpm. 200 nM S-CDK were added to DDK-phosphorylated MCM-DHs, dCMGE assembly was then started by further addition of 40 nM Dpb11, 32.5 nM Pol epsilon, 133 nM GINS, 106 nM Cdc45, 40 nM Sld3/7, and 67 nM Sld2. After 12 minutes of incubation at 30°C and 1,250 rpm, the reactions were immediately used for negative stain grid preparation. Wild-type proteins were substituted with mutant constructs or dropped out as specified. To split dCMGEs into single CMGEs (sCMGEs), 100 nM Mcm10 and 300 nM RPA were added simultaneously with the other firing factors.

### In vitro pre-IC assembly

To assemble the pre-IC, DDK-phosphorylated MCM-DHs were prepared as described above, but eluted in the absence of ATP. Sld3/7 and Sld2 were replaced by equimolar amounts of ppSld3/7 and phosphomimetic Sld2^8D^, and no CDK was added. DDK-phosphorylated DHs were incubated with ppSld3/7, Sld2^8D^, Dpb11, Pol epsilon, GINS and Cdc45 in the absence of ATP and CDK for 10 minutes at 24°C and 1,250 rpm, after which reactions were either negatively stained, applied to cryo-EM grids and plunge frozen using a Vitrobot, or further purified.

### ALFA pulldown of pre-IC and dCMGE complexes

To purify either pre-IC or dCMGE, phospho-DH formation and respective complex assembly was carried out on an ALFA-tagged 2xMH-ARS1 DNA template. 20 µL of assembly reaction prepared as described above was added to 15 µL ALFA PE Selector slurry (NanoTag Biotechnologies) equilibrated in Buffer P and bound for 5 minutes at 24°C and 1,250 rpm.

For pre-IC maturation reactions, beads were washed with 3x 150 µL Buffer P and eluted for 5 minutes at 24°C and 1,250 rpm in 20 µL Buffer P + 200 µM ALFA peptide (NanoTag Biotechnologies). Pre-IC to dCMGE conversion was triggered by adding 1 mM ATP per reaction and analysed by negative-stain EM.

To assay for high-salt stability of pre-IC and dCMGE, beads were washed with 3x 150 µL of Buffer R (25 mM HEPES-KOH pH 7.5, 5 mM Mg(OAc)_2_, 10% v/v glycerol, 0.02% w/v NP-40) + either 250 mM K-Glutamate (Low-Salt Wash) or 300 mM KCl (High-Salt Wash). Elution was carried out in 20 µL Buffer P + 200 µM ALFA peptide (no nucleotide for pre-IC reactions, 1 mM ATP for dCMGE reactions) for 5 minutes at 24°C and 1,250 rpm.

### In vitro DNA replication assay

DNA replication was performed at 30°C and 1250 rpm following previous protocols^74^. In short, MCM-DHs were loaded for 20 minutes using 40 nM ORC, 40 nM Cdc6, 60 nM Mcm2-7/Cdt1, 4 nM 10.6kb pJY22 plasmid template DNA in Buffer S (25 mM HEPES-KOH pH 7.6, 100 mM potassium glutamate, 10 mM magnesium acetate, 5 mM ATP, 0.02% NP-40-S and 2 mM DTT). Loaded MCMs were phosphorylated with 50 nM DDK for 15 min. DNA replication was performed for 30 min by adding final concentrations of 40 nM Dpb11, 20 nM Pol epsilon, 20 nM GINS, 80 nM Cdc45, 20 nM CDK, 25 nM Sld3/7, 50 nM Sld2, 200 nM RPA, 20 nM TopoI, 50 nM Pol alpha, 20 nM Mcm10, 200 μM each CTP/GTP/UTP, 80 μM each dNTP and 33 nM C^32^P-dCTP. When specified, Dpb11, Sld3/7 and Sld2 proteins were excluded from reactions or substituted by mutant versions. In the ‘CDK Bypass experiment’, reactions contained either 10 nM WT Sld2 and 5 nM Sld3/7 or 50 nM Sld2-8D and 25 nM phosphorylated Sld3/7.

Reactions were stopped with 85 mM EDTA, cleared over an Illustra MicroSpin G-50 column, denatured in 2% sucrose/0.02% bromophenol blue/60 mM NaOH/10 mM EDTA, separated on 0.8% agarose gels in alkaline conditions containing 30 mM NaOH and 2 mM EDTA at approximately 1 volt/cm for 17h, fixed in cold 5% trichloroacetic acid, dried, exposed to phosphor screens, and scanned using a Typhoon phosphor imager.

### Sample preparation and data collection for nsEM

Carbon-coated 300-mesh copper grids (EM Resolutions) were glow-discharged at 25 mA for 1 minute in a GloQube Plus (Quorum) in ambient air. 4 µL sample were incubated for 2 minutes on a glow-discharged grid, blotted, and negatively stained by two applications of 4 µL 2% Uranyl-Acetate for 20 seconds, after which the grid was blotted dry. NS-EM micrographs were acquired using a Rio16 camera (Gatan Digital Micrograph) on a FEI Tecnai G2 Spirit Twin microscope operated at 120 KV. Approximately 50 – 150 micrographs were collected per dataset at 3.1 Å/px (29,000 x magnification) at -1 to -2 µm defocus.

### nsEM image processing

Micrographs were imported in Relion 4^75^, Contrast Transfer Function (CTF) was estimated using Gctf^76^. MCM-containing particles were picked using crYOLO 1.9.2^77^, imported into Relion and extracted with a 144 px box size. Extracted particles were 2D-classified in either Relion 4 or cryoSPARC v4.4.1^78^. Interpretable class averages were categorized (DH, pre-IC, cis- or trans-dCMGE, sCMGE) and particles were quantified. Complex assembly efficiency was calculated as number of all detected target particles divided by the number of all licensed replication origins. For example, number of pre-IC particles were divided by the sum of DHs and pre-IC particles. For Mcm10-dependent CMGE splitting experiments, all sCMGEs were multiplied by a factor of 0.5, to account for the fact that sCMGEs originate from a single, licensed replication origin.

### Cryo-EM sample preparation of pre-IC

MCM-DHs were loaded onto an ALFA-tagged 2xMH-ARS1 template and phosphorylated by DDK as described above. 100 µL DNA-loaded, phospho-DHs were purified on 15 µL ALFA PE Selector beads for 10 minutes at 24°C and 1,250 rpm. Subsequently, beads were washed 3x with 200 µL Buffer Q and 1x with 200 µL Buffer O and eluted for 10 minutes in 20 µL Buffer O supplemented with 200 µM ALFA peptide at 24°C and 1250 rpm. Given that ALFA elution yielded a higher number of phospho-DHs compared to StrepTactinXT purification, the eluted phospho-DHs were incubated with 4x higher molar amount of firing factors (Dpb11, Pol epsilon, GINS, Cdc45, ppSld3/7, Sld2^8D^) at 24°C and 1,250 rpm for 10 minutes. Graphene-oxide coated 300-mesh UltrAuFoil R1.2/R1.3 grids were prepared on the day following a previously published protocol^47^.4 µL sample were applied per grid in a Mark IV Vitrobot (FEI) and incubated for 60 seconds at 24°C and 90% humidity. Grids were blotted for 3 – 4.5 seconds at blotforce 0 and plunge-frozen in liquid ethane.

### Cryo-EM data collection of pre-IC

59,347 movies were collected on a 300 kV FEI Titan Krios G3i at a nominal magnification of 130,000 x (0.95 Å/px physical pixel size) using a Falcon 4 direct electron detector (DED) in counting mode and a Selectris energy filter with a 10 eV slit width using EPU v3.2. Three shots were acquired per hole at spot size 9 with a beam diameter of 660 nm, a 100-µm objective aperture inserted and a defocus range from -2.0 to -3.0 µm. Each movie was recorded with 1674 Electron Event Representation (EER) frames for 5.44 seconds with a total fluence of 39 electrons/Å^2^ (**Extended Data Table 1**).

### Cryo-EM image processing of pre-IC

59,347 EER movies were aligned and dose-weighted with 5 x 5 patches using Relion’s own implementation of MOTIONCOR2^79^. 54 internal frames were grouped into 31 fractions resulting in a dose per frame of 1.26 electrons/Å^2^. Motion-corrected micrographs were imported into cryoSPARC v4.4.1^78^, CTF estimation was carried out via Patch CTF. Initial particle picking was performed using Blob Picker with a diameter range from 200 – 350 Å and circular blobs. 5,149,760 particles were extracted fourfold binned to 3.8 Å/px with a 150 px box size and cleaned up with multiple rounds of reference-free 2D classification with 400 classes and an uncertainty factor of 2. 62,192 MCM double hexamer and pre-initiation complex particles were selected from 2,867 micrographs containing more than 20 particles per micrograph in a defocus range from -1.5 to -2.5 µm and used to train a Topaz network with 75 expected particles per micrograph^80^. Following Topaz picking at an extraction threshold of -6 with an extraction radius of 28 px, 1,572,464 particles were extracted and Fourier-cropped to 3.8 Å/px with a box size of 150 px and subjected to three rounds of 2D classification yielding 302,557 double hexamers and 345,093 pre-initiation complexes.

Ab-initio reconstructions of both DHs and preICs were generated independently in C1, the initial volumes of each complex were used as 3D references for heterogenous refinement to further separate DHs and preICs from each other and remove low-quality particles. 279,730 DH particles were separated into 20 classes by alignment-free 3D classification in cryoSPARC. 162,764 high-quality DH particles were selected, unbinned to 0.95 Å/px and refined to a final resolution of 2.8 Å with C2 symmetry applied. After separating pre-IC particles from DH particles through heterogenous refinement, 335,896 binned pre-IC particles were NU-refined with C2 symmetry applied, unbinned to 0.95 Å/px with a 600 px box size and 3D-classified without alignment yielding 290,496 particles that were NU-refined to 3.72 Å. This first unbinned reconstruction of the pre-IC was subjected to further classification in cryoSPARC to remove low-quality particles, resulting in a stack of 151,175 pre-IC particles displaying improved density quality. Following homogenous refinement, symmetry expansion was performed. Local C1 refinement yielded a 3.4 Å resolution map of the pre-IC dimer, with some residual anisotropy visible. To improve the reconstruction further, refinement of a signal subtracted monomer was performed. To generate the best possible mask for subtraction, a double-subtraction strategy was implemented. First one asymmetric unit was locally refined using a soft mask around the top pre-IC monomer. The resulting, improved, reconstruction was used to generate a new soft mask for signal subtraction, used to accurately remove the signal from the top pre-IC monomer. The remaining pre-IC monomer was reconstructed and locally refined to 3.3 Å resolution, featuring improved isotropy. This map was used to generate a third soft mask for a second signal subtraction of the bottom monomer. This allowed us to determine the structure of the top monomer using local refinement to a resolution of 3.2 Å.

### Cryo-EM sample preparation of phospho-DH-3745

DDK-phosphorylated, DNA-loaded MCM-DHs were eluted from MagStrep “type3” XT beads in Buffer T (25 mM HEPES-KOH pH 7.5, 100 mM KOAc, 0.02% w/v NP-40, 25 mM D-Biotin) and incubated with 32 nM (yeast-expressed) Sld3/7 and 127 nM Cdc45 for 10 minutes at 30°C and 1,250 rpm. To stabilise the phospho-DH3745 complex, the sample was crosslinked with 0.05% w/v glutaraldehyde for 5 minutes and quenched with 25 mM Tris-HCl pH 7.5. For the first dataset, the reaction was vitrified without further purification as described for the pre-IC. For the second and third dataset, the reaction was purified after crosslinking via ALFA pulldown. 5 crosslinked and quenched reactions were pooled and bound to 1.5 µL ALFA Selector PE resin through the ALFA-tagged 2xMH-ARS1 DNA template for 1 hour at 24°C and 1,250 rpm, washed once with 50 µL Buffer O and eluted for 30 minutes at 24°C and 1,250 rpm in 12 µL Buffer O + 200 µM ALFA peptide. Cryo-EM grids were prepared as described for the pre-IC.

### Cryo-EM data collection of phospho-DH-3745

Three datasets of 31,794 (#1), 32,200 (#2), and 41,658 (#3) movies were acquired on a 300 kV FEI Titan Krios G3i using a Gatan K2 Summit direct electron detector in counting mode and a BioQuantum energy filter with a slit width of 20 eV at a nominal magnification of 130,000 x (1.08 Å/px) using EPU v3.2. Per hole, 2 shots were recorded with a 100-µm objective aperture inserted, a defocus range from -1.1 to -2.5 µm and a total fluence of 49.1 – 50.4 electron/Å^2^.

### Cryo-EM image processing of phospho-DH-3745

Each dataset was pre-processed separately. First, cryo-EM movies were motion-corrected in Relion 4 using its own implementation of MOTIONCOR2^79^, CTF estimation was carried out with Gctf^76^ A Topaz picking network *^80^* was iteratively trained on the first dataset using a selection threshold of -3, a scale factor of 8 and 30 expected particles per micrograph. Picked particles were extracted 2x binned (2.16 Å/px) with a 360-px box size and cleaned up by reference-free 2D classification. Noise and low-quality averages were discarded. Remaining particles were used as input for the next round of Topaz training. Ultimately, particles were unbinned (448-px box size) and transferred to cryoSPARC^78^ to generate an initial volume and subjected to non-uniform refinement with C2 symmetry applied. 100,495 particles from dataset 1, 184,975 particles from dataset 2 and 239,120 particles from dataset 3 were then joined for downstream processing, yielding 524,590 particles. After another round of 2D classification, 359,025 particles underwent 2 rounds of CTF-refinement (beam-tilt, anisotropic magnification, per-particle defocus and per-micrograph astigmatism) and Bayesian polishing in Relion*^81^,* resulting in a 3.1 Å reconstruction of the consensus phospho-DH bound to Sld3 after non-uniform refinement in cryoSPARC with C2 symmetry applied.

To isolate Cdc45-bound phospho-DHs, particles were first C2-symmetry expanded in cryoSPARC. An AlphaFold multimer^82^ prediction of a Sld3 CBD-Cdc45 complex was aligned with the map of the phospho-DH by docking an atomic model of a CMG ring (PDB 7QHS)^30^ into one MCM ring and superimposing the prediction via Cdc45. A volume of the aligned Sld3 CBD-Cdc45 prediction was generated at 10 Å resolution through the “molmap” command in ChimeraX^83^ and used as input to prepare a soft mask in Relion. Using this mask, 718,050 C2 symmetry-expanded phospho-DH particles were subjected to two rounds of focused 3D classification without alignment and a T-value of 20 into 8 classes. 72,693 particles displaying proteinaceous features inside this mask were selected and refined in Relion in C1 local searches restricted to 1.8° while masking out Sld3 CBD-Cdc45 signal from the opposite MCM hexamer. Subsequently, the masked out Sld3 CBD-Cdc45 density was signal subtracted in Relion. The Cdc45-bound DH was then imported into cryoSPARC and locally refined, yielding a final reconstruction at 3.7 Å.

In parallel, C2-symmetry expanded Sld3-bound phospho-DHs were subjected to signal subtraction within cryoSPARC. Cdc45 and Sld3 CBD density associated with one of the two MCM hexamers was masked out and subtracted. 718,050 particles of the remaining phospho-DH bound to a single Sld3 CBD-Cdc45 complex were subjected to two rounds of 3D classification without alignment in cryoSPARC. Ten classes were chosen, with a class similarity of 0.25 and resolution was limited to 15 Å. 96,027 particles were selected, from 3D classes displaying featured Cdc45 density, and locally refined using C1 symmetry using cryoSPARC to a resolution of 3.5 Å. This approach yielded a map of the Sld3-Cdc45-bound phospho-DH, also featuring Sld7 density.

### Cryo-EM sample preparation of sCMGE assembled with Sld2 and RPA

Single CMGE complexes on double-roadblocked ARS1 DNA (168 bp) were essentially prepared as described above (In vitro ATP-dCMGE assembly). Following dCMGE splitting, 20-µL assembly reactions were further purified on paramagnetic ALFA beads as described before (ALFA pulldown of pre-IC and dCMGE complexes). Cryo-EM grids were prepared as described for the pre-IC, with 3 applications of 4 µL eluate per grid.

### Cryo-EM data collection of sCMGE assembled with Sld2 and RPA

50,060 movies were collected at a nominal magnification of 130,000 x (0.95 Å/px physical pixel size) on a FEI Titan Krios G3i using a FalconIV direct electron detector (DED) in counting mode and a Selectris energy filter with a 10 eV slit width using EPU v3.2. Per hole, three shots were acquired with a defocus range from -2.0 to -2.9 µm and a 100-µm objective aperture inserted. Movies were recorded with 31 frames and a total dose of 38.6 electrons/Å^2^ (**Extended Data Table 2**).

### Cryo-EM image processing of sCMGE assembled with Sld2 and RPA

50,060 EER movies were motion-corrected using Relion’s own implementation of MOTIONCOR2^79^ and CTF-estimated with CTFFind 4.1.13^84^. 1,940 particles were manually picked from 20 micrographs and used as input for Topaz training^80^. Using Topaz, 4,827,716 particles were picked and extracted at 3.8 Å/px (4x binning) and a 108 px box size, and cleaned up with multiple rounds of 2D Classification in cryoSPARC v4.4.1^78^. A subset of clean sCMGE and MCM-DH classes were selected to generate initial 3D models, which were used in Heterogenous Refinement of a total of 3,801,543 sCMGE and MCM-DH particles. This yielded 1,125,095 sCMGE particles, which were unbinned with a 432 px box and homogeneously refined (with global CTF correction) to a final resolution of 2.7 Å.

### Cryo-EM sample preparation of sCMGE assembled without Sld2 and RPA

Single CMGE complexes were assembled as described above with one notable exception, omission of Sld2. After dCMGE splitting, 13 20-µL reactions were pooled and jointly purified on 60 µL ALFA beads. Binding and Washing steps were carried out as described, and DNA-bound complexes eluted in 50 µL Buffer O + 200 µM ALFA Peptide. Lacey cryo-EM grids (400 mesh Cu; TAAB) were coated with Graphene oxide by first hydrophilising the grid surface with 4 µL 300 nM DDM and side-blotting, followed by two rounds of on-grid incubation with 4 µL of a 20 µg/mL Graphene oxide suspension in 300 nM DDM. Grids were washed with 3 5-µL droplets of mH2O from the backside, and blotted dry from the backside. Cryo-EM grids were vitrified as described before for the pre-IC, with 2 applications of 4 µL eluate per grid.

### Cryo-EM data collection of sCMGE assembled without Sld2 and RPA

70,337 movies were recorded on a FEI Titan Krios G3i at a nominal magnification of 130,000 x (0.95 Å/px physical pixel size) using a Falcon IV DED in counting mode and a Selectris energy filter with a 10 eV slit width using EPU v3.2. Shots were acquired in a 0.7 µm spacing, with a 100-µm objective aperture inserted and a defocus range from -1.4 to -2.4 µm. Per movie, 29 frames were recorded with a total fluence of 42.0 electrons/Å^2^ (**Extended Data Table 2**).

### Cryo-EM image processing of sCMGE assembled without Sld2 and RPA

70,337 EER movies were preprocessed using Relion’s own implementation of MOTIONCOR2^79^ and imported into cryoSPARC v4.4.1^78^ for CTF estimation via PatchCTF. 4,579 particles were manually picked from 134 micrographs and used to train a Topaz model^85^ with 75 expected particles per micrograph. 2,390,783 particles were extracted with 8x binning at 7.6 Å/px and a 70 px box size, and subsequently cleaned up with multiple rounds of reference-free 2D classification, removing well-averaging MCM-DHs as well as noise, ultimately yielding 162,691 CMG-like particles. Initial 3D references were generated with clean subsets of sCMGE- and dCMGE-like classes (orange and purple brackets, respectively, **Extended Data Fig. 8c**) using cryoSPARCs Ab initio reconstruction. These volumes were subsequently used to separate sCMGEs from contaminating dCMGE-like particles through multiple rounds of Heterogenous refinements (C1 symmetry, 1 sCMGE reference, two dCMGE references). 72,370 cleaned-up sCMGE particles were then unbinned in Relion 5.0 with a box size of 512 px, submitted to 2 rounds of Bayesian Polishing and 3 rounds of CTF Refinement (each round correcting separately: 4th order aberrations, tilt and trefoil; anisotropic magnification; per-particle defocus and per-micrograph astigmatism) and 3D-refined using Blush^86^ to a nominal resolution of 3.4 Å.

### Atomic model building and refinement

The pre-IC complex structure was built to a locally-refined monomeric pre-IC cryo-EM map density-modified with EMReady^87^. A single CMGE complex extracted from PDB entry 7PMK^48^ was initially docked into the cryo-EM density. After this initial placement, individual domains were then docked as rigid bodies into the cryo-EM density using UCSF Chimera^88^. As they strikingly matched the cryo-EM density, the structure predictions of Dpb11-Mcm7, Dpb11-GINS, Cdc45-Sld3, Mcm4-Sld3 and Mcm7-Sld7 generated with AlphaFold3^37^ were also used as a starting point for modelling these interaction interfaces. Each chain or pair of chains were flexibly fit after generating self-restraints in Coot^89^ using density maps of varying blurring. Fragments mapping outside the visible density were deleted. The entire model was then manually adjusted with real-space refinement in Coot,, using varying whole molecule restraints depending on the local quality of the density. Automated real space refinement was then performed in Phenix version 1.21^90^ against the non-postprocessed map.

The pre-IC dimer complex structure was built to a C2-symmetric cryo-EM map density-modified with EMReady. First, two refined pre-IC monomers were rigid-body docked into the cryo-EM density map in ChimeraX. DNA chains from each monomer were trimmed at the overlapping region and were merged. Finally, protein-protein interface between the monomers was adjusted using real space refinement in Coot. Automate real space refinement was then perfomred in Phenix version 1.21^90^ against the non-postprocessed map.

The DH-Sld3-MBD and DH-3745 complex was built starting from a previously published MCM double hexamer structure bound to duplex DNA (PDB entry 7P30)^14^, combined with the structure predictions of Mcm4-Sld3, Sld7-Mcm6 and Cdc45-Sld3 produced using AlphaFold2 v2.3.4^82^. Initial docking was performed in ChimeraX^91^. Density fit of protein regions except zinc fingers was first adjusted using molecular-dynamics based real space refinement in Isolde^92^. Next, positions of all atoms were adjusted with flexible fitting and real space refinement (sphere refinement) in Coot^89^. Fragments mapping outside the visible density were truncated from the atomic coordinate file. Automated real space refinement against the non-postprocessed map.

The structure of sCMGE assembled on ARS1 DNA with RPA and Sld2 was modeled into a cryo-EM map density-modified with EMReady, using PDB 7PMK^48^ as an initial template. The starting model was rigid-body docked in UCSF Chimera, followed by flexible fitting of individual chains in Coot using chain restraints. Where density permitted, the model was expanded manually or guided by AlphaFold2 predictions. Due to insufficient resolution for base identification, an arbitrary repetitive DNA sequence was modelled. To account for the bases that stretch between the modelled dsDNA and ssDNA stretches and could not be built due to poor density, we maintained the nucleotide numbering from PDB entry 6SKL^93^. Final automated real-space refinement was performed using Phenix (v1.21) against the non-postprocessed map. For all structures, the quality of the resulting atomic models was evaluated with MolProbity^94^ (**Extended Data Tables 1 and 2**). Structures in figures were displayed using EMReady-postprocessed maps.

## General

We would like to thank members of the Costa lab for useful discussion, the Crick Structural Biology STP for computational support (A. Purkiss and A. Nans); yeast cultures (N. Patel, A. Alidoust and D. Patel) and cryo-EM support (A. Nans and N. Lukyanova).

## Author contributions

T.P. and A.C. conceived the study. T.P. performed all wet lab and structure determination work. Exceptions are the sCMGE(+Sld2) preparation and structure done by G.P. and the DNA replication assays, done by B.C. supervised by J.F.X.D. O.W. and

J.S.L. and contributed to cloning, expression and purification. J.S.L. also developed ALFA-tag purification with T.P. A.B. and E.C.C. contributed to structure determination. T.P. and A.C. wrote the paper with input from all other authors. A.C. supervised the study.

## Data availability

Data supporting the findings of this study are available within the paper and its supplementary information files. Cryo-EM density maps have been deposited in the Electron Microscopy Data Bank (EMDB) under the following accession codes: EMD-53972 (maps for the DH-Sld3-MBD), EMD-53973 (maps for the DH-Sld3/7-Cdc45), EMD-53971 (maps for the dimeric pre-IC), EMD-53970 (maps for the monomeric pre-IC), EMDB-56898 (maps for sCMGE assembled with Sld2 and RPA), EMD-56897 (maps for sCMGE assembled with RPA and without Sld2). Atomic coordinates have been deposited in the Protein Data Bank (PDB) with the following accession codes: 9RHL (DH-Sld3-MBD), 9RHM (DH-Sld3/7-Cdc45), 9RHJ (dimeric pre-IC), 9RHI (monomeric pre-IC) and PDB 28VY (sCMGE assembled with Sld2 and RPA).

## Extended Data Figure Legends

**Extended Data Figure 1.**
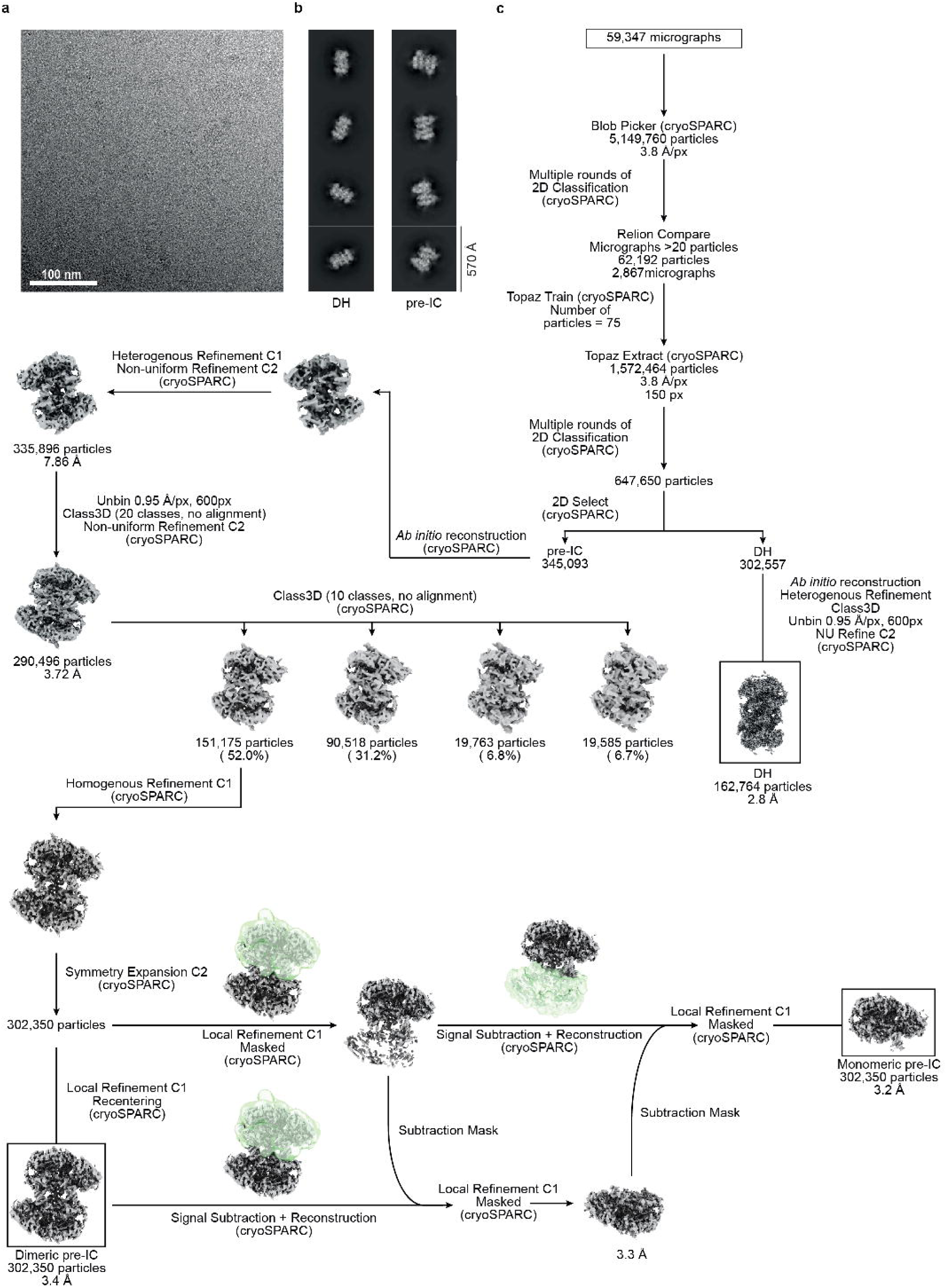
Cryo-EM imaging and image processing workflow for the pre-IC. **(a)** Representative cryo-electron micrograph. This cryo-EM sample was prepared and imaged more than three times. **(b)** Cryo-EM 2D averages of the DH and pre-IC assembly. **(c)** Image processing pipeline.

**Extended Data Figure 2.**
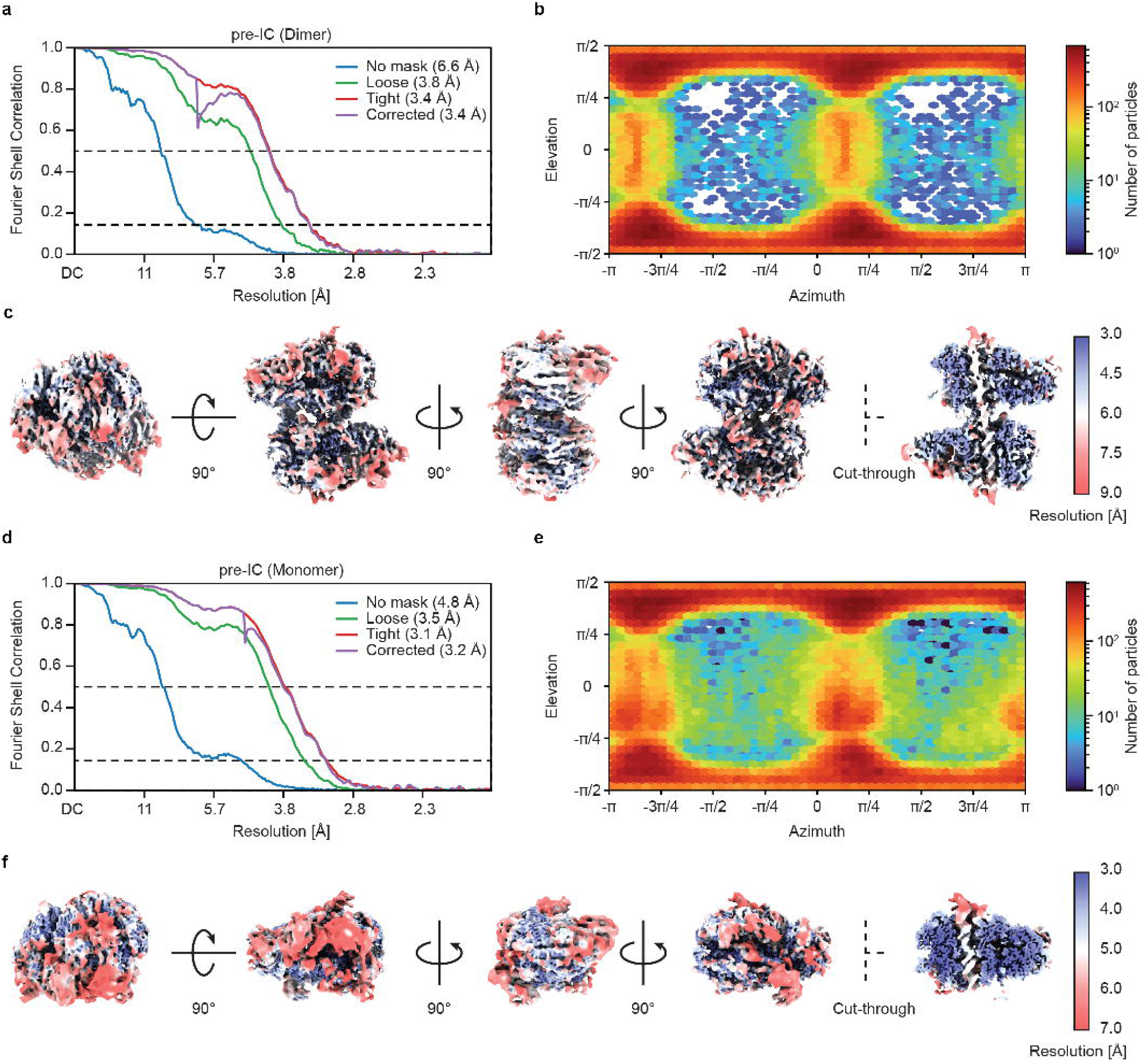
Validation of the pre-IC map. **(a)** Fourier shell correlation of the full dimeric pre-IC map. **(b)** Angular distribution plot of the full dimeric pre-IC map. **(c)** Five views of the of the full dimeric pre-IC map colored according to the local resolution. **(d)** Fourier shell correlation of the signal-subtracted, monomeric pre-IC map. **(e)** Angular distribution plot of the signal-subtracted, monomeric pre-IC map. **(f)** Five views of the of the signal-subtracted, monomeric pre-IC map colored according to the local resolution.

**Extended Data Figure 3.**
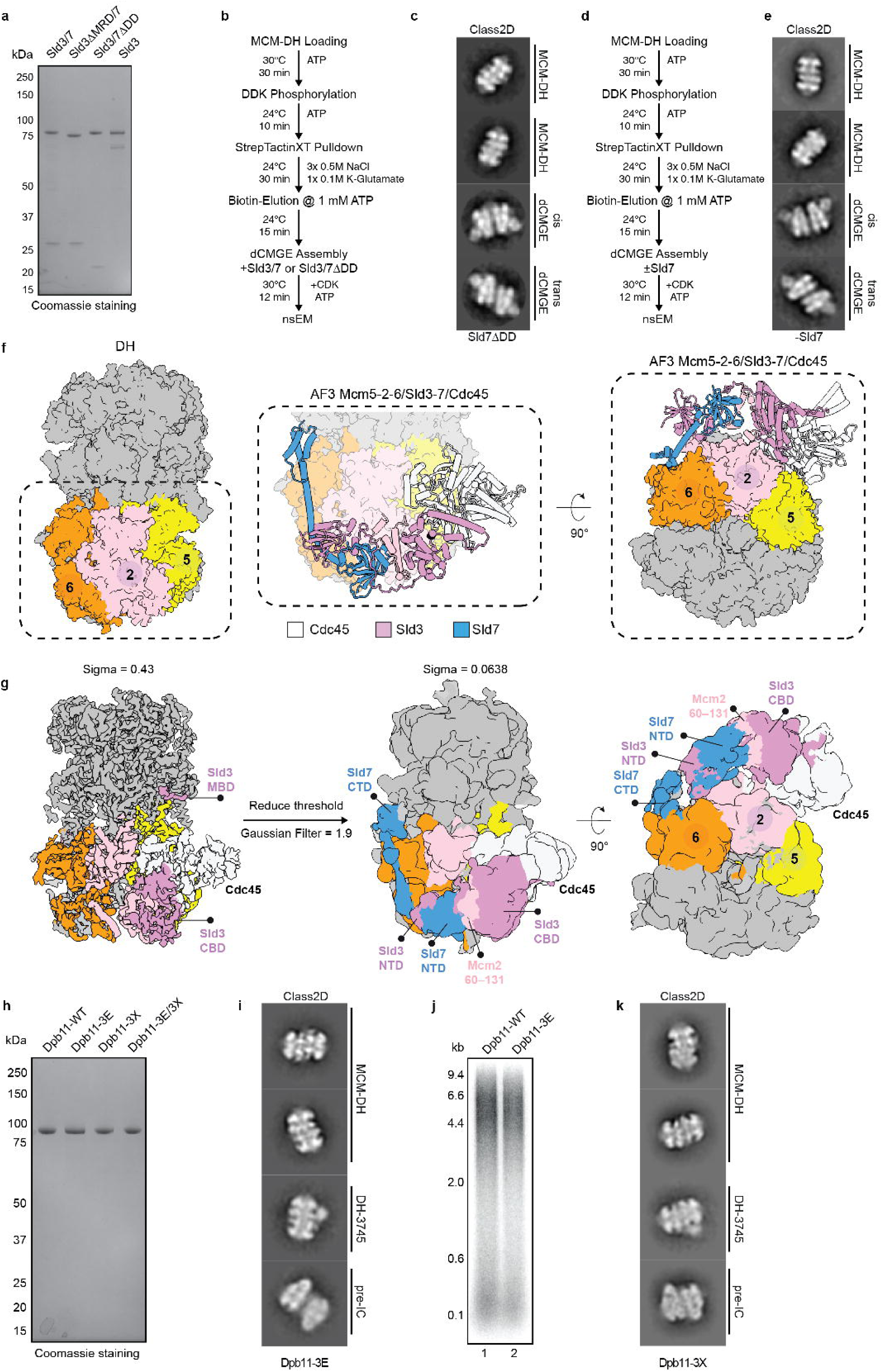
Sld7 requirement, Sld3/7-Cdc45 architecture on the DH, and Pre-IC assembly with Dpb11 variants. **(a)** SDS PAGE of purified Sld3 constructs. This gel was reproduced three times. **(b-c)** Workflow and representative 2D classes for pre-IC assembly with wild type or Sld7 DD truncation mutant. **(d-e)** Workflow and representative 2D classes for dCMGE formation with or without Sld7. **(f)** AF3 modelling of the entire Sld3-Sld7-Cdc45 interaction with Mcm6,2,5, overlayed with the experimentally determined DH structure. **(g)** A cryoSPARC derived volume of the DH-3745 Gaussian-filtered in ChimeraX displaying the network of interactions between Sld3-Sld7 and Cdc45 on the DH. **(h)** Coomassie stained gel for the wild type and mutant Dpb11 targeting the Mcm7 (3E), GINS (3X) or both (3E/3X) binding sites. This gel was repeated twice. **(i)** 2D averages for the pre-IC formation reaction assembled with Dpb11-3E. **(j)** DNA replication reactions with wild type and 3E variant of Dpb11. A small defect for the variant protein can be detected. The bigger effect observed in pre-IC assembly by EM (Fig. 4d) indicates that assessing the formation of this transient structural intermediate is a more sensitive assay compared to DNA replication reconstituted in vitro. This experiment was performed two times. **(k)** 2D averages for the pre-IC formation reaction assembled with the Dpb11-3X mutant. For gel source data, see Supplementary Fig. 1.

**Extended Data Figure 4.**
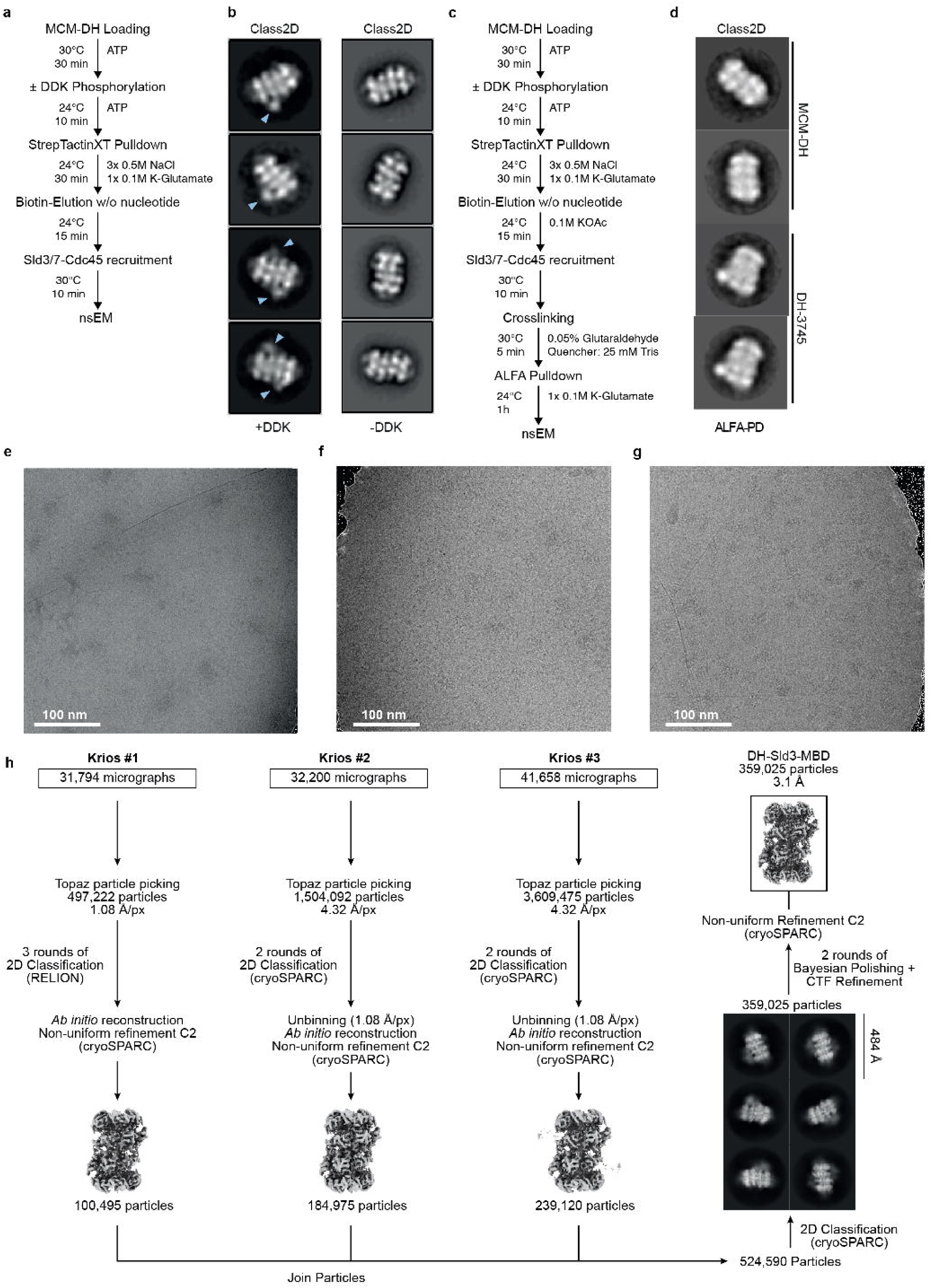
DH-Sld3/7-Cdc45 assembly and processing pipeline for the DH-Sld3-MBD. **(a)** Workflow for the preparation of the non-crosslinked DH-Sld3/7-Cdc45. **(b)** 2D averages obtained by supplementing (left) or omitting (right) DDK. The blue arrows indicate Cdc45 density. Our first nsEM attempts resulted in decoration of DHs with poorly averaged density on the side of either one or both MCM hexamers. **(c)** Workflow for the preparation of DH-Sld3/7-Cdc45 (crosslinked and purified). **(d)** 2D averages obtained for the crosslinked and purified DH-Sld3/7-Cdc45. Improved DH decoration was obtained upon glutaraldehyde crosslinking followed by nanobody-affinity purification using an ALFA tag**^95^**, a second, crosslinking-tolerant handle fused to the MH DNA roadblock). **(e-g)** Representative cryo-electron micrographs from each individual dataset. The respective samples were reconstituted more than three times and imaged under cryo conditions two (EDF 4e) and four (EDF 4f-g) times. **(h)** Pipeline for determining the DH-Sld3-MBD structure.

**Extended Data Figure 5.**
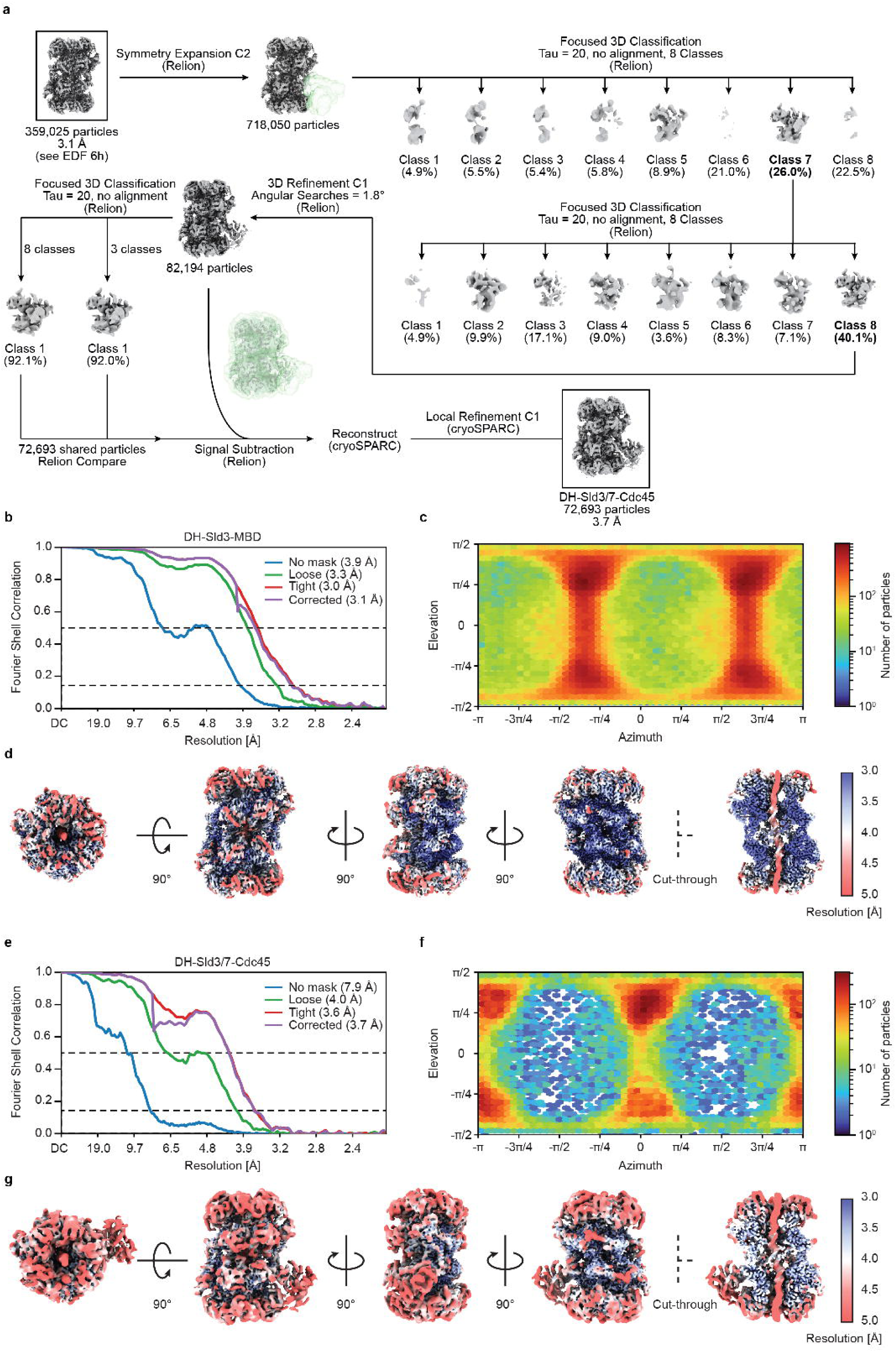
Processing pipeline for DH-Sld3/7-Cdc45 and validation of the DH-Sld3-MBD and DH-Sld3/7-Cdc45 maps. **(a)** Pipeline for determining the DH-Sld3/7-Cdc45 structure. **(b)** Fourier shell correlation of the DH-Sld3-MBD map. **(c)** Angular distribution plot of the full dimeric DH-Sld3-MBD map. **(d)** Five views of the of the DH-Sld3-MBD map colored according to the local resolution. **(e)** Fourier shell correlation of the DH-Sld3/7-Cdc45 map. **(f)** Angular distribution plot of the DH-Sld3/7-Cdc45 map. **(g)** Five views of the of the DH-Sld3/7-Cdc45 map colored according to the local resolution.

**Extended Data Figure 6.**
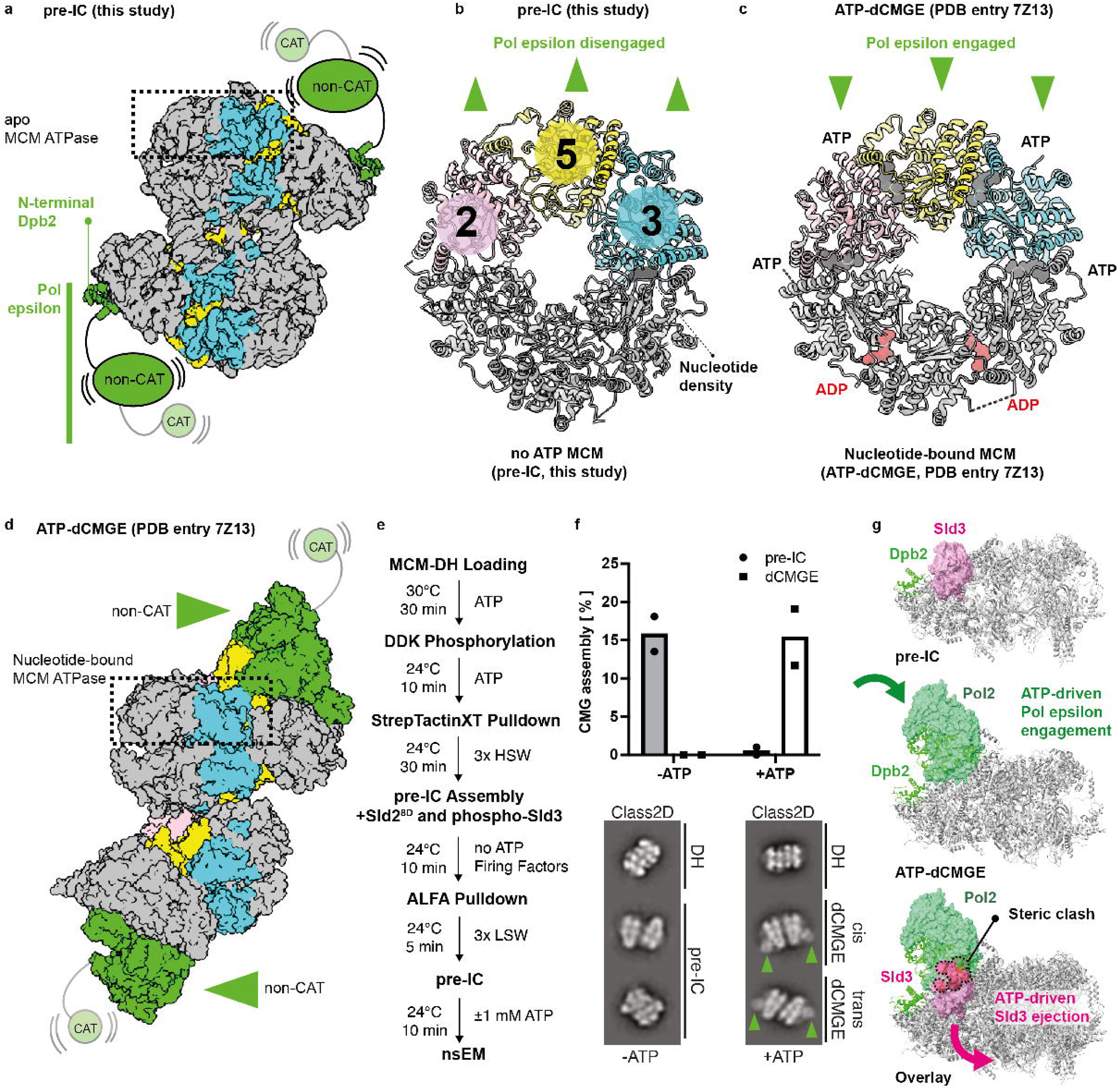
Pol epsilon in the pre-IC. **(a)** The only Pol epsilon element visible in the pre-IC is N-terminal Dpb2. **(b)** In the pre-IC, no nucleotide density can be observed within the Mcm2-5 interface. In the same structure, Pol epsilon is disengaged from the ATPase tier of MCM. **(c)** In the ATP-dCMGE structure, the Mcm2-5 interface is ATP bound and Pol epsilon engages the ATPase domain. **(d)** Side view of ATP-dCMGE, with the non-catalytic Dpb2-C-terminal-Pol2 portion of Pol epsilon visible. **(e)** Flow chart of pre-IC formation in apo conditions, followed by addition of ATP. **(f)** In apo conditions only pre-IC complexes and no dCMGE can be observed. When ATP is spiked in, the majority of preIC complexes are converted to dCMGE, where Pol epsilon decorates CMG on the ATPase face of MCM. This experiment was performed two times. **(g)** Mechanism for Sld3 eviction, through the ATP-dependent engagement of C-terminal Pol2 to the ATPase side of MCM.

**Extended Data Figure 7.**
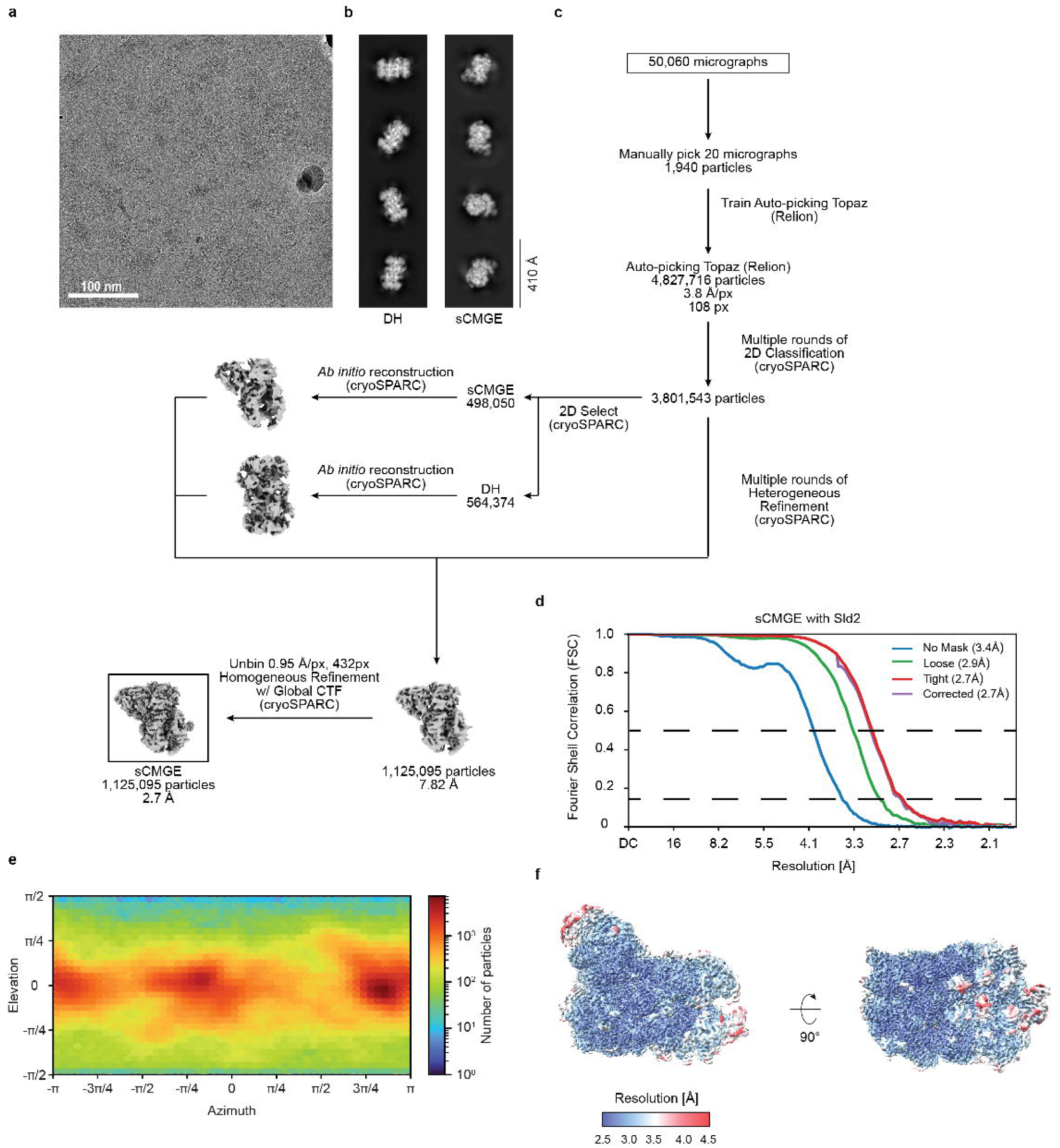
Processing pipeline and validation of the sCMGE assembled at the ARS1 origin with Mcm10, RPA and Sld2. **(a)** Representative cryo-electron micrograph. This sCMGE formation experiment was repeated three times. **(b)** 2D averages of inactive DHs and activated sCMGEs. **(c)** Image processing pipeline. **(d)** Fourier shell correlation plot. **(e)** Angular distribution. **(f)** sCMGE filtered and colored according to the local resolution.

**Extended Data Figure 8.**
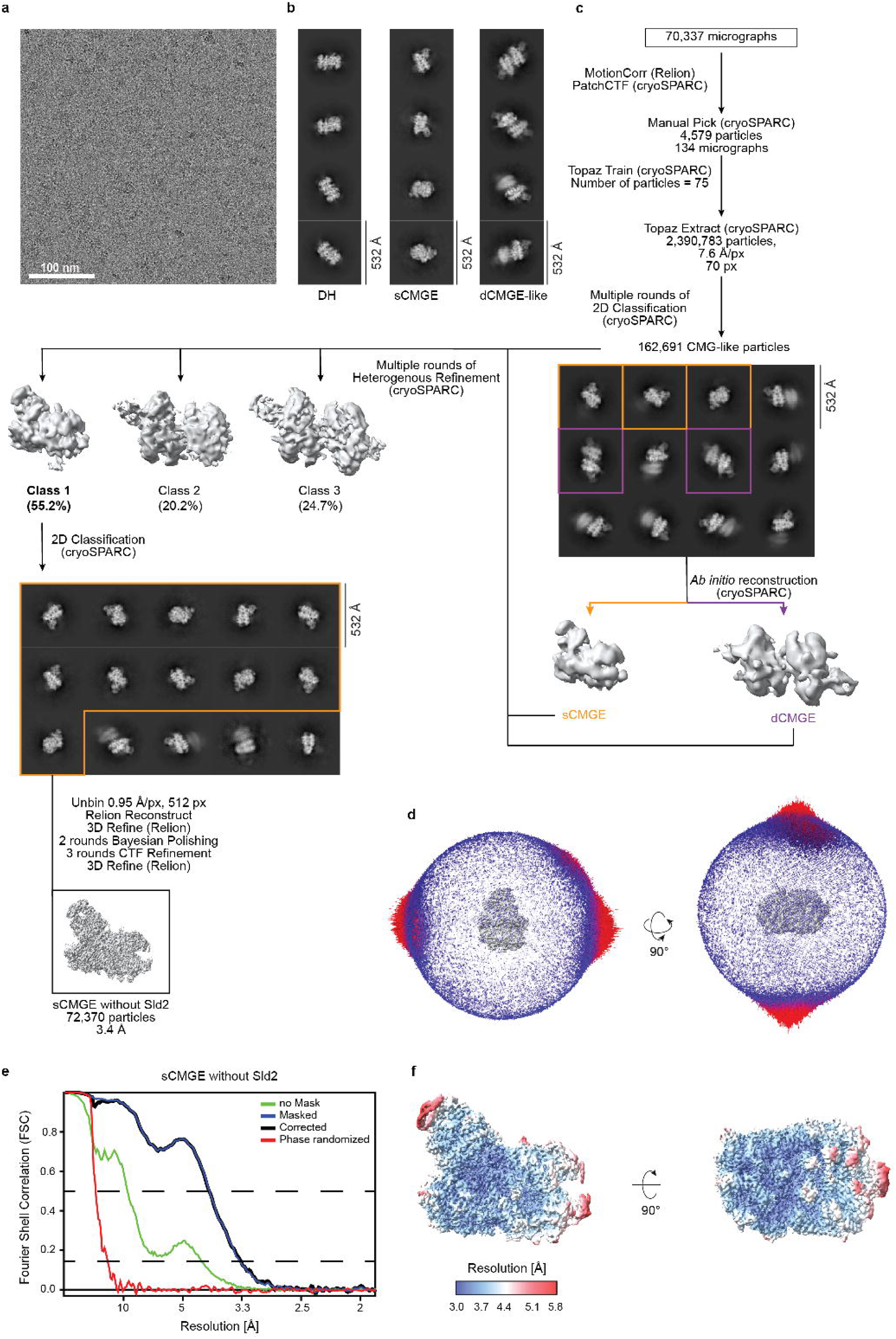
Processing pipeline and validation of the sCMGE assembled at the ARS1 origin with Mcm10, RPA and without Sld2. **(a)** Representative cryo-electron micrograph. This sCMGE formation experiment was repeated three times. **(b)** 2D averages of inactive DHs, sCMGEs and split sCMGEs. **(c)** Image processing pipeline. **(d)** Angular distribution. **(e)** Fourier shell correlation plot. **(f)** sCMGE filtered and colored according to the local resolution.

**Extended Data Table 1. Cryo-EM data collection, refinement and validation statistics.** Information for the structures of Pre-IC Monomer, Pre-IC Dimer, DH-Sld3-MBD, DH-3745 is presented.

**Extended Data Table 2. Additional cryo-EM data collection, refinement and validation statistics.** Information for the structures of sCMGE assembled with or without Sld2 in the presence of RPA is presented.

