## Supplemental Information for "Structure of the Pre-Initiation Complex Explains CMGE Biogenesis"

### 1. Supplementary Results

#### Modulation of Sld3 binding by Rad53

Rad53 halts origin firing by targeting Sld3, as well as the Dbf4 subunit of DDK[1]. In a short flexible insertion bracketed by two alpha helices built in our atomic model we could identify three known Rad53 target residues (T519, S521 and T524)[1,2], located immediately adjacent the A4-engaged element of Sld3 (**Fig. 2e** and **Supplementary Fig. 5a**). This observation was striking given that four DDK target residues in Mcm4 (T140, S141, S144, S145)[3] and four Rad53 target residues in Dbf4 (S518, S521, S526, S528)[1] also map within an A4-binding element, in the non-phosphorylated DH and DH-DDK complex respectively (**Supplementary Figs. 5b,c**). This suggests that regulation of MCM binding via phosphorylation by two distinct kinases has evolved to target three proteins that bind the same Mcm4 epitope (Mcm4 itself, Dbf4 and Sld3), which engage the helicase motor sequentially, on the path to origin activation.

To test this model, we used AF3[4] to predict the effect of phosphorylation on these three Mcm4 interactions. In our positive control, AF3 recapitulated the phospho-dependent disengagement of the N-terminal Mcm4 from A4 in cis, previously described with cryo-EM[3,5,6] (**Supplementary Fig. 5b**). AF3 then predicted that phosphorylation of Rad53 target residues in Dbf4 would abrogate A4 engagement (**Supplementary Fig. 5c**). This agrees with our previous nsEM analysis showing that Rad53-phosphorylated DDK fails to engage DHs[3]. Finally, AF3 predicted that phosphorylation of Rad53 target residues in Sld3 abrogates A4 engagement (**Supplementary Fig. 5a**). This prediction implies that impairing the A4-Sld3 interaction should prevent Cdc45 recruitment to MCM. To test this, we performed dCMGE assembly with unmodified or Rad53 pre-phosphorylated Sld3/7. While dCMGE formed efficiently without Rad53[7], Rad53-prephosphorylated, re-purified Sld3/7 blocked its assembly (**Supplementary Figs. 6a-d**). Our data agree with previous observations that Sld3 phosphorylation by Rad53 blocks Cdc45 recruitment to MCM[1,8].

#### The Dpb11 phospho-reader function

Dpb11 plays an essential phospho-reader function, however the local resolution of the BRCT1-2 does not allow building phosphorylation-dependent inter-subunit contacts. Despite these limitations, superposition of the co-crystal structure of *S. pombe* orthologs (Rad4 bound to phospho-Sld3[9]) reveals that the phospho-Sld3 binding sites on Dpb11 are solvent-accessible in the pre-IC structure. This result is also recapitulated through AlphaFold modelling showing that *S. cerevisiae* Sld3 phospho-Thr600 and phospho-Ser622, essential for replication initiation, engage the BRCT1 and 2, as previously described with genetics and biochemical analysis[10,11]. When modelled in the full context of the pre-IC experimental structure we note that the C-terminal end of the Sld3 atomic model points towards, and maps close to phospho-Thr600 (64 Å apart, **Supplementary Figs. 7a-d**). This measurement suggests a model whereby Sld3 physically engages Mcm4-6-7 and Dpb11 concurrently.

#### Changes in DNA engagement upon DH-3745 to pre-IC transition

Disrupting DH dimerisation upon pre-IC formation involves a substantial structural transition, with a change in buried surface area from 6,816 to 722 Å$^2$. What might provide the energy for this reconfiguration? Several changes in the nucleoprotein assembly suggest a mechanism, although the sequence of events remains unclear. For example, comparison with the DH reveals that ADP is released upon pre-IC formation (**Supplementary Figs. 3a,b**), which could promote long-range allosteric changes. Also, MCM binding by GINS-Cdc45 appears to rearrange the N- and C-terminal tiers of MCM[12,13], likely contributing to the disruption of the DH dimerisation interface. The most striking structural change, however, occurs for the DNA double helix, which transitions from a compressed to a relaxed state, similar to a loaded spring snapping free. Within DH and DH-3745, in fact, the two MCM rings are slightly offset so that duplex DNA must bend to traverse the entire length of the central channel[3,14,15]. This high-energy state is relieved upon pre-IC formation, as the double helix reverts to its basal (straight) B-form state (**Supplementary Fig. 8a**). This transition is rendered possible by a change in the way ATPase pore loops grip DNA. In the DH, helix 2 insert (h2i) loops arrange in a planar configuration. Upon pre-IC formation h2i loops let go of DNA and become arranged in a left-handed coil around DNA, resulting in an anti-parallel helical structure. This is different from the right-handed helical bias observed for ATP-bound CMGE, which promotes DNA melting and unwinding downstream of pre-IC formation (**Supplementary Fig. 8b**).

### 2. Supplementary Figures

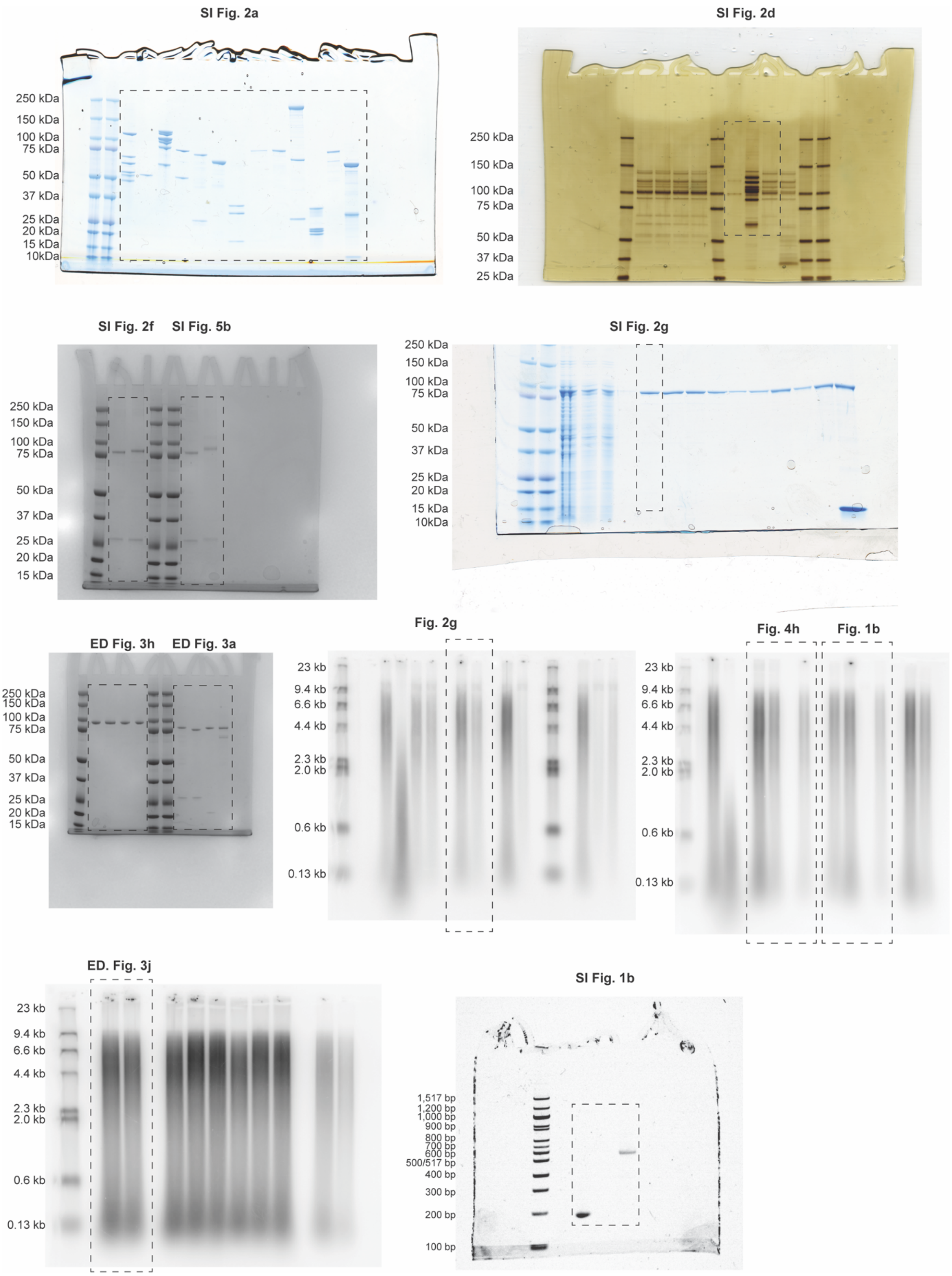


**Supplementary Figure 1 | Gel Source Data.** Uncropped SDS PAGE, EMSA and DNA replication assay images. Dashed line boxes indicate the cropped area shown in the respective display items.

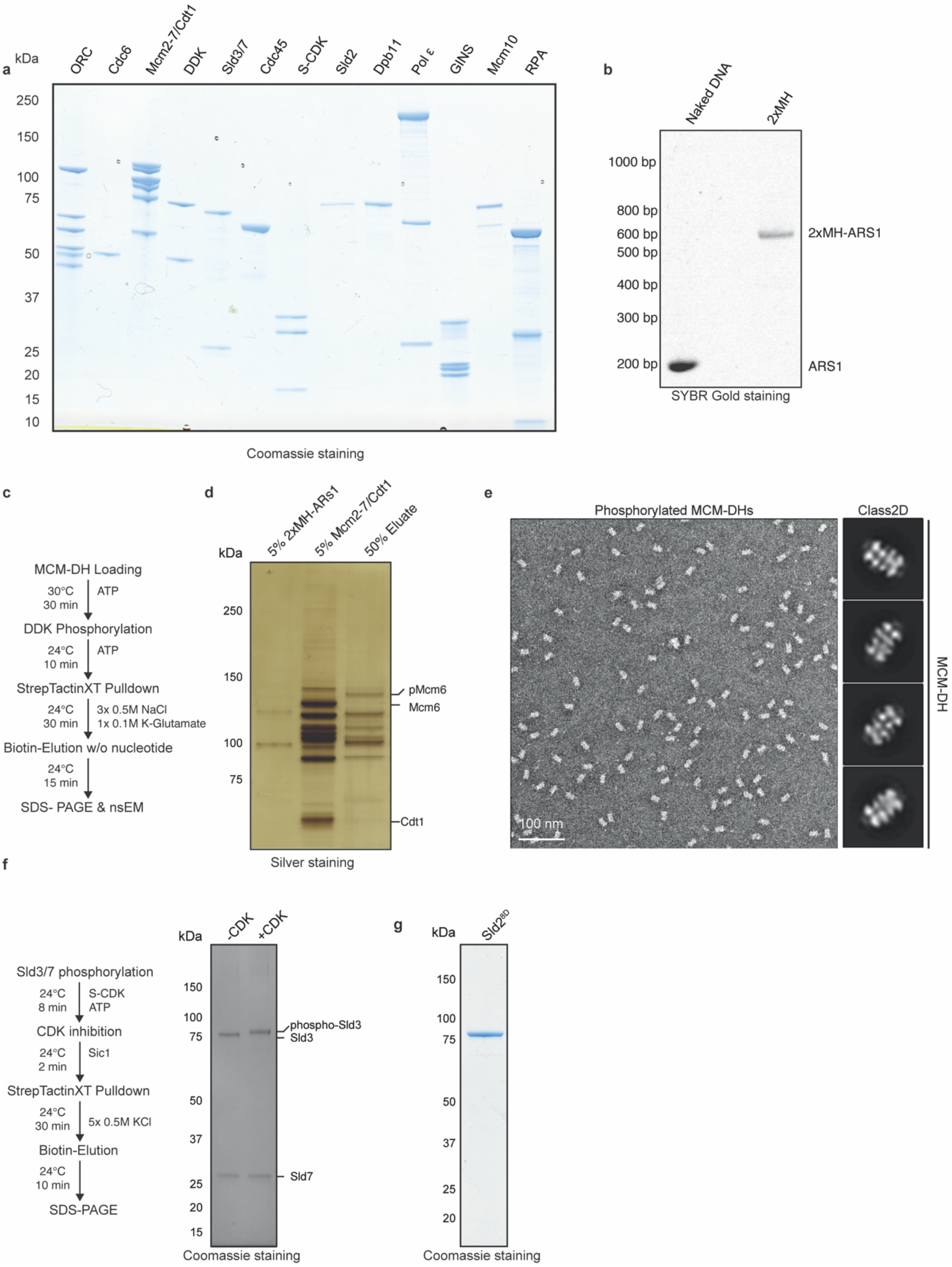


**Supplementary Figure 2 | Reagents used for assembling the pre-IC. (a)** Purified proteins. This result was reproduced more than three times. **(b)** ARS1 origin of replication blocked at both ends with HpaII methyltransferase (MH). This experiment was repeated more than three times. **(c)** Workflow for producing DNA-loaded, DDK-phosphorylated DHs. **(d)** Silver-stained gel of the MH-blocked DNA, the loading-competent MCM-Cdt1 complex and eluted DNA-loaded phospho-DHs. This assay was replicated more than three times. **(e)** Negative stain micrograph and 2D averages of the purified phospho-DHs. This imaging experiment was carried out more than three times. **(f)** On the left, workflow for producing pre-phosphorylated Sld3/7 using S-CDK. On the right, Coomassie-stained gel of the non-phosphorylated and phosphorylated Sld3/7 complex. This experiment was repeated three times. **(g)** Purified phospho-mimicking Sld2$^{8D}$ variant. This purification was repeated three times. For gel source data, see Supplementary Fig. 1.

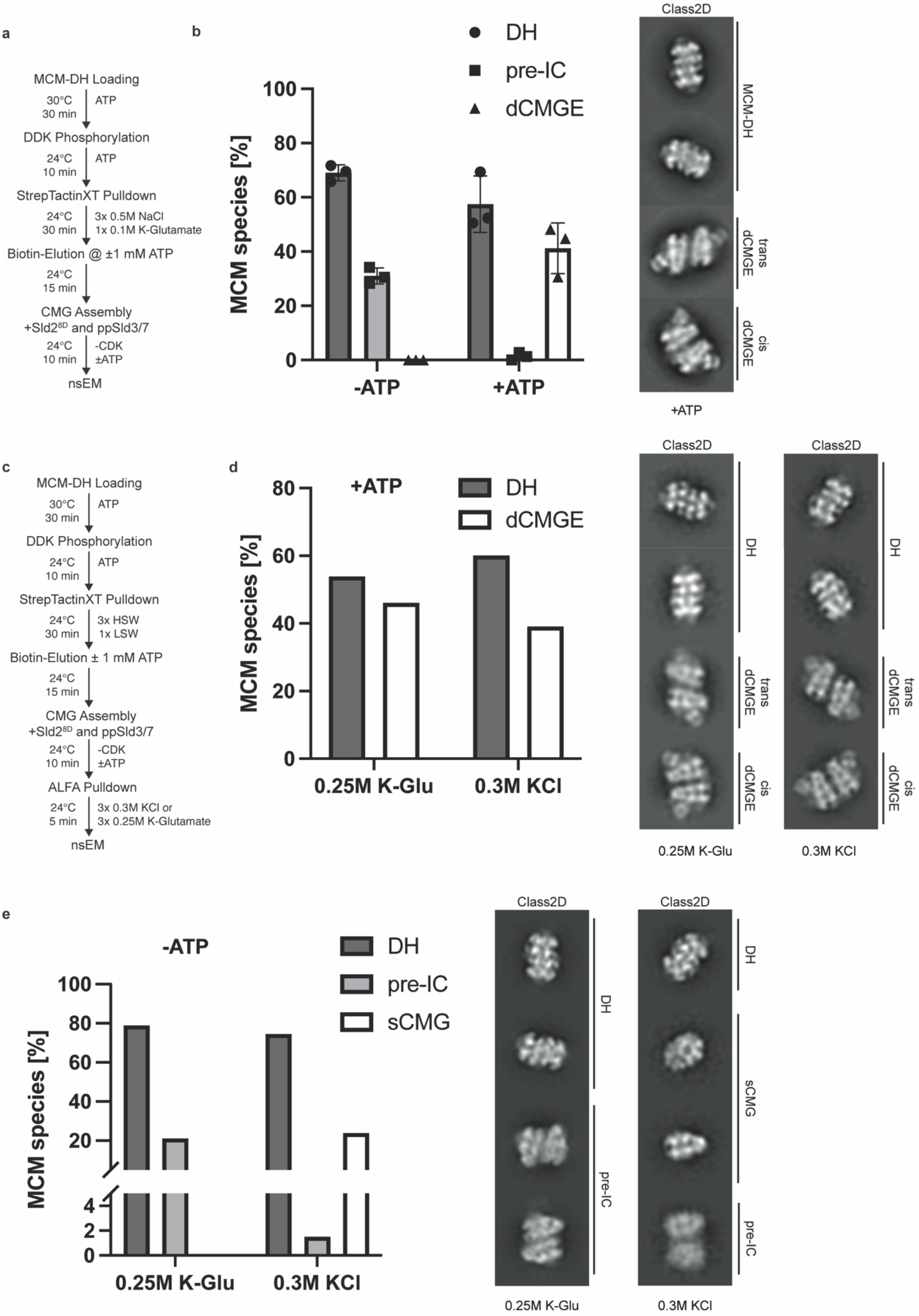
a
MCM-DH Loading
30°C 30 min ATP
DDK Phosphorylation
24°C 10 min ATP
StrepTactinXT Pulldown
24°C 30 min 3x 0.5M NaCl 1x 0.1M K-Glutamate
Biotin-Elution @ ±1 mM ATP
24°C 15 min
CMG Assembly
+Sld2^8D and ppSld3/7
24°C 10 min -CDK ±ATP
nsEM
b
DH
pre-IC
dCMGE
MCM species [%]
-ATP
+ATP
Class2D
MCM-DH
trans dCMGE
cis dCMGE
+ATP
c
MCM-DH Loading
30°C 30 min ATP
DDK Phosphorylation
24°C 10 min ATP
StrepTactinXT Pulldown
24°C 30 min 3x HSW 1x LSW
Biotin-Elution ± 1 mM ATP
24°C 15 min
CMG Assembly
+Sld2^8D and ppSld3/7
24°C 10 min -CDK ±ATP
ALFA Pulldown
24°C 5 min 3x 0.3M KCl or 3x 0.25M K-Glutamate
nsEM
d
+ATP
DH
dCMGE
MCM species [%]
0.25M K-Glu
0.3M KCl
Class2D
DH
trans dCMGE
cis dCMGE
0.25M K-Glu
0.3M KCl
e
-ATP
DH
pre-IC
sCMG
MCM species [%]
0.25M K-Glu
0.3M KCl
Class2D
DH
sCMG
pre-IC
0.25M K-Glu
0.3M KCl

**Supplementary Figure 3 | Assembly and negative stain 2D averages of dCMGE and pre-IC.** **(a)** Workflow of dCMGE formation in the absence of CDK. **(b)** On the left, bar graph of DH-to-pre-IC conversion efficiency in the absence of ATP and DH-to-dCMGE conversion efficiency in the presence of ATP. On the right, negative stain 2D averages of DHs and dCMGEs in *trans* and *cis* configurations. 2D averages of the no ATP condition are displayed in Fig. 1c. Error bars, mean ± s.d. This experiment was performed three times. **(c)** Workflow for high-salt stability challenge of pre-IC and dCMGE. **(d)** On the left, dCMGE and DHs (+ATP assembly) challenged with different salt washes. On the right, representative 2D averages of complexes imaged after salt challenge. **(e)** On the left, pre-IC and DHs (-ATP assembly) challenged with different salt washes. On the right, representative 2D averages of complexes imaged after salt challenge. Some pre-IC complexes disassemble into single CMGs.

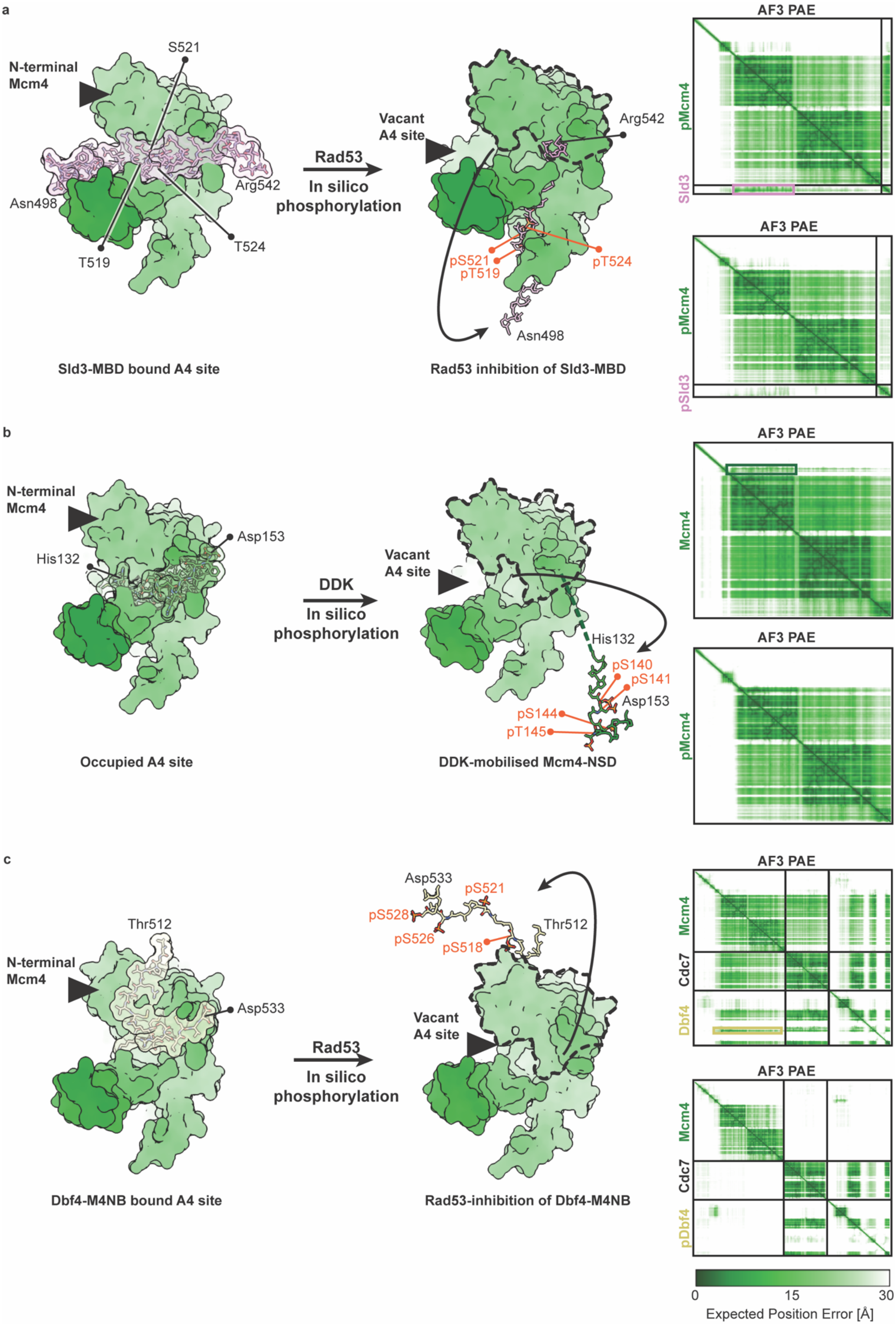
a
N-terminal Mcm4
S521
Asn498
Arg542
T519
T524
Rad53
In silico phosphorylation
Vacant A4 site
Arg542
pS521
pT519
pT524
Asn498
Sld3-MBD bound A4 site
Rad53 inhibition of Sld3-MBD
AF3 PAE
pMcm4
Sld3
AF3 PAE
pMcm4
pSld3
b
N-terminal Mcm4
Asp153
His132
DDK
In silico phosphorylation
Vacant A4 site
His132
pS140
pS141
Asp153
pS144
pT145
Occupied A4 site
DDK-mobilised Mcm4-NSD
AF3 PAE
Mcm4
AF3 PAE
pMcm4
c
Thr512
N-terminal Mcm4
Asp533
Rad53
In silico phosphorylation
Asp533
pS521
pS528
pS526
pS518
Thr512
Vacant A4 site
Dbf4-M4NB bound A4 site
Rad53-inhibition of Dbf4-M4NB
AF3 PAE
Mcm4
Cdc7
Dbf4
AF3 PAE
Mcm4
Cdc7
pDbf4
0
15
30
Expected Position Error [Å]

**Supplementary Figure 4 | AlphaFold 3 predictions of Mcm4 binding by client proteins regulated through phosphorylation. (a)** On the left, unphosphorylated Mcm4. In the center, Mcm4 with DDK target sites phosphorylated disengages the A domain. On the right, Predicted Alignment Error (PAE) plots for the two structure predictions. **(b)** On the left, unphosphorylated Dbf4 subunit of DDK engages the Mcm4 A domain. The Cdc7 subunit of DDK is not shown. In the centre, Dbf4 with Rad53 target sites phosphorylated disengages Mcm4. On the right, Predicted Alignment Error (PAE) plots for the two structure predictions. **(c)** On the left, unphosphorylated Sld3 engages the Mcm4 A domain. In the centre, Sld3 with Rad53 target sites phosphorylated disengages the A domain. On the right, Predicted Alignment Error (PAE) plots for the two structure predictions.

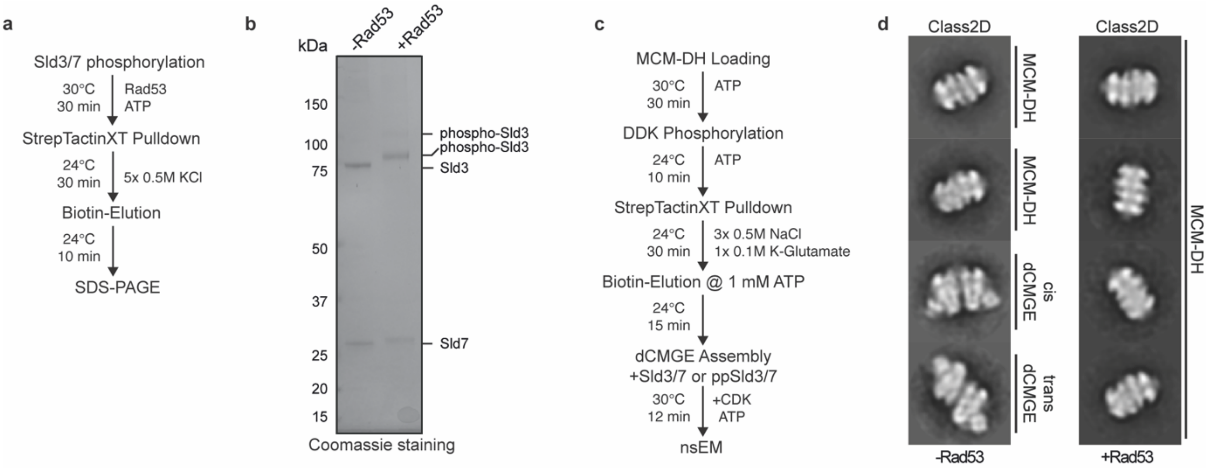


**Supplementary Figure 5 | Sld3 phosphorylation by Rad53.** **(a)** Workflow for Sld3/7 pre-phosphorylation by Rad53. **(b)** SDS PAGE of untreated and Rad53-treated Sld3/7 after affinity purification. For gel source data, see Supplementary Fig. 1. **(c)** Workflow of dCMGE formation experiment comparing untreated and Rad53-treated Sld3/7. This experiment was performed three times **(d)** Representative nsEM 2D averages of the experiment described in (c).

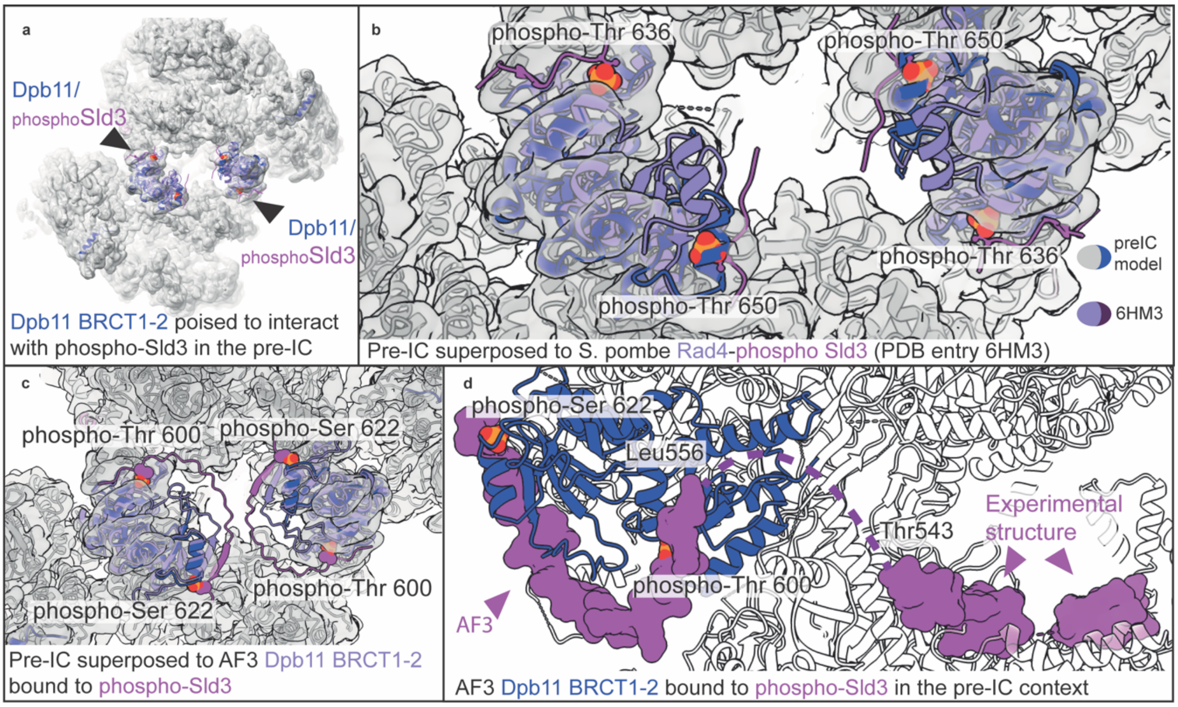


**Supplementary Figure 6 | Dpb11 phosphoreader function. (a)** Phospho-Sld3 binding sites on Dpb11 BRCT1-2 are solvent-accessible in the pre-IC structure. **(b)** Superposition of the S. pombe Rad4-phospho-Sld3 co-crystal structure (PDB entry 6HM3) to the Dpb11 BRCT1-2 domains in our S. cerevisiae pre-IC structure. **(c)** AF3 prediction of Dpb11 BRCT1-2 bound to phospho-Sld3 superposed to our pre-IC experimental structure. **(d)** The C-terminal end of Sld3 in our experimental structure is perfectly poised for joining with phospho-Thr600 of the AF3 prediction.

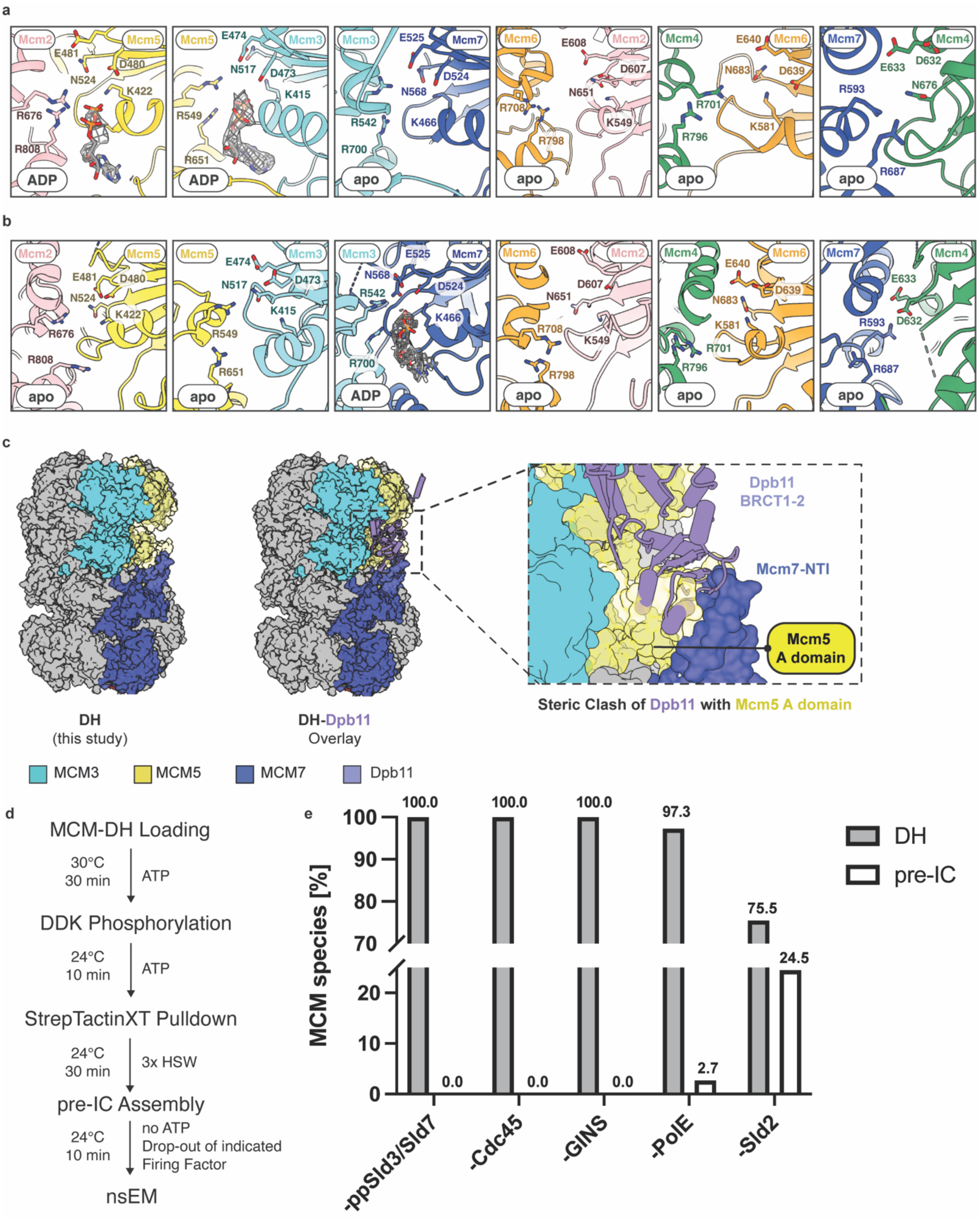


**Supplementary Figure 7 | Nucleotide occupancy, duplex DNA trajectory and steric clashes in DH-Sld3/7-Cdc45 and pre-IC, and Pre-IC assembly in the absence of individual firing factors. (a)** Nucleotide occupancy at the six ATPase sites of the DH-Sld3/7-Cdc45 structure. Compared to the previously published DH in ATP[14], one difference is in the Mcm6-2 ATPase site, which features loosely-interacting ATPase subunits compatible with a released-nucleotide state. Overall nucleotide occupancy is also different, with ADP retained only at the Mcm2-5 and Mcm5-3 sites. **(b)** Nucleotide occupancy at the six ATPase sites of the pre-IC structure. **(c)** Dpb11 engagement of Mcm7, as observed in the pre-IC, is incompatible with DH formation. In fact, Dpb11 clashes with Mcm5. **(d)** To establish what role Pol epsilon has in CMG formation, we first assembled pre-IC with or without Pol epsilon. We found that Pol epsilon stimulated but was not strictly required for pre-IC formation (though efficiency was dropped dramatically in Pol epsilon dropout conditions). This argues for an ancillary role of Pol epsilon in GINS recruitment, supported by previous observations that GINS forms a pre-loading complex with Pol epsilon, Dpb11 and phospho-Sld2, aiding the CMG formation process[16]. Workflow for pre-IC assembly while dropping out selected firing factors. **(e)** Bar graph displaying the fraction of DHs and pre-IC complexes in the absence of ppSld3/7, Cdc45, GINS, Pol epsilon or Sld2.

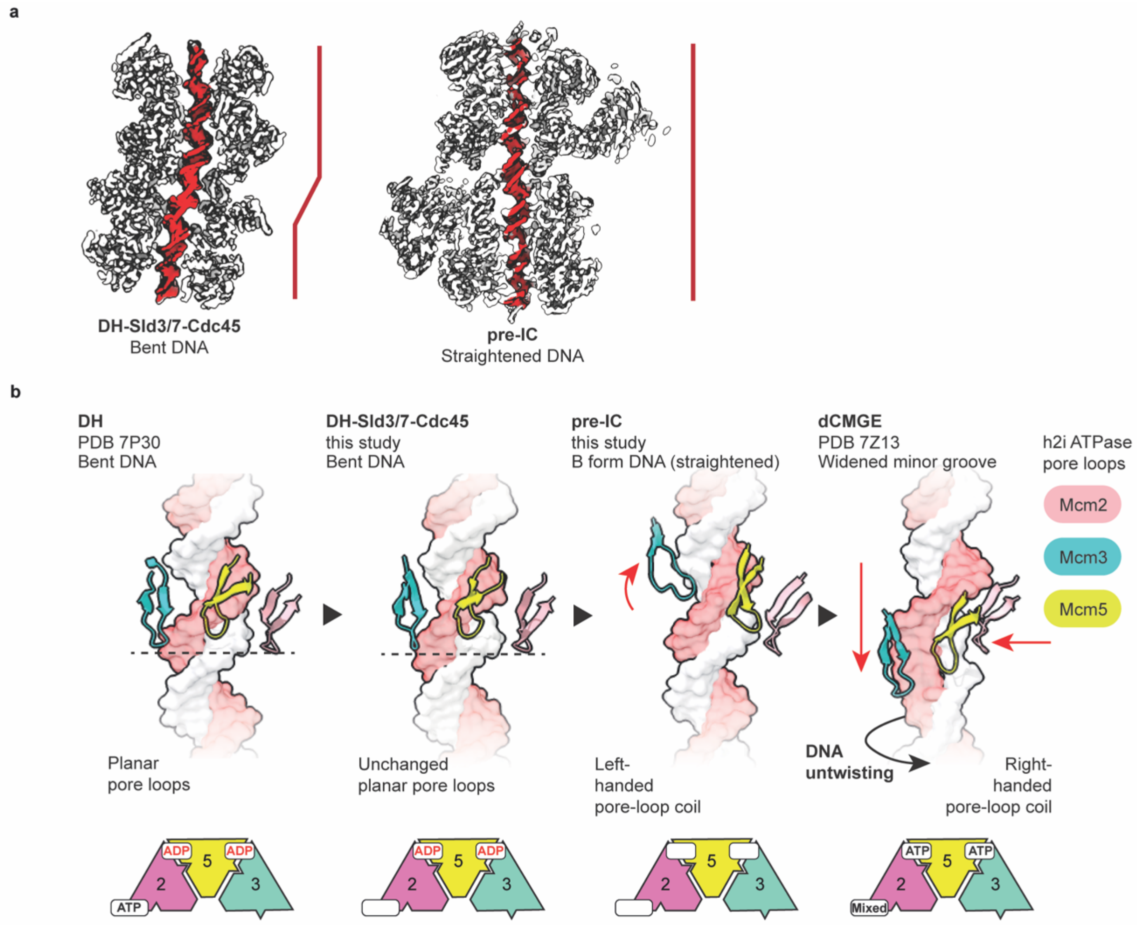


**Supplementary Figure 8 | Changes in DNA engagement during CMGE biogenesis. (a)** Cut-through view of the DH-Sld3/7-Cdc45 and pre-IC structures highlights different trajectory for the DNA double helix. In the first structure (left), DNA bends as it runs through the central channel at the interface between the two cockeyed MCM rings. In the pre-IC, where the MCM dimerisation interface is reconfigured, DNA runs straight. **(b)**Transition between the DH to the DH-Sld3/7-Cdc45 to the pre-IC to the dCMGE structure involves nucleotide release and rebinding and a rearrangement of the h2i ATPase pore loops. In the ADP/ATP loaded DH, the Mcm2,3,5 pore loop are arranged in a planar configuration. This configuration remains unchanged when nucleotide is released by Mcm2 (upon DH-Sld3/7-Cdc45 complex formation. The same h2i pore loops form a left-handed coil in the pre-IC structure, upon further nucleotide release. ATP binding causes assembly factor ejection and maturation to the double CMGE, which untwists and melts DNA, in preparation for the opening of a replication bubble.

### 3. Supplementary Tables

| Name | Use | Composition |
|---|---|---|
| Buffer A | Protein Purification: Sld3/7, Dpb11 | 25 mM HEPES-KOH pH 7.5, 500 mM KCl, 10% v/v Glycerol, 0.02% w/v NP-40, 1 mM EDTA, 1 mM DTT |
| Buffer B | Protein Purification: Dpb11 | 25 mM HEPES-KOH pH 7.5, 150 mM KCl, 10% v/v Glycerol, 0.02% w/v NP-40, 1 mM EDTA, 1 mM DTT |
| Buffer C | Protein Purification: Dpb11 | 25 mM HEPES-KOH pH 7.5, 300 mM KOAc, 10% v/v Glycerol, 0.02% w/v NP-40, 1 mM EDTA, 1 mM DTT |
| Buffer D | Protein Purification: Sld2 | 25 mM HEPES-KOH pH 7.5, 800 mM KCl, 10% v/v Glycerol, 1M Sorbitol, 2 mM ATP, 10 mM $MgCl_2$, 0.02% v/v NP-40, 0.1% w/v Tween-20, 1 mM DTT |
| Buffer E | Protein Purification: Sld2 | 25 mM HEPES-KOH pH 7.5, 500 mM NaCl, 10% v/v Glycerol, 0.02% w/v NP-40, 1 mM EDTA, 1 mM DTT |
| Buffer F | Protein Purification: Sld2 | 25 mM HEPES-KOH pH 7.5, 700 mM KOAc, 40% v/v Glycerol, 0.02% w/v NP-40, 1 mM EDTA, 1 mM DTT |
| Buffer G | Protein Purification: Pol epsilon | 25 mM HEPES-KOH pH 7.6, 400 mM KOAc, 10% v/v Glycerol, 2 mM DTT |
| Buffer H | Protein Purification: Sic1 | 25 mM HEPES-KOH pH 7.5, 500 mM NaCl, 10% v/v Glycerol, 1 mM EGTA, 0.2% w/v Triton X-100, 0.5 mM TCEP, 10 mM Imidazole |
| Buffer I | Protein Purification: Sic1 | 25 mM HEPES-KOH pH 7.5, 5% v/v Glycerol, 5 mM $MgCl_2$, 0.5 mM EDTA, 0.5 mM TCEP |
| Buffer J | Protein Purification: Mcm10 | 25 mM HEPES-KOH pH 7.6, 500 mM NaCl, 10% v/v Glycerol, 1 mM EDTA, 0.05% w/v Tween-20, 1 mM DTT |
| Buffer K | Protein Purification: Mcm10 | 25 mM HEPES-KOH pH 7.6, 300 mM NaCl, 10% v/v Glycerol, 0.05% w/v Tween-20, 1 mM DTT, 5 mM Desthiobiotin |
| Buffer L | Protein Purification: Mcm10 | 25 mM HEPES-KOH pH 7.6, 200 mM NaCl, 20% v/v Glycerol, 0.05% w/v Tween-20, 1 mM EDTA, 2 mM DTT |
| Buffer M | Sld3/7 Prephosphorylation | 40 mM HEPES-KOH pH 7.5, 310 mM K-Glutamate, 10 mM $Mg(OAc)_2$, 10% v/v Glycerol, 0.02% w/v NP-40, 1 mM DTT, 2 mM ATP, 0.4 mg/mL BSA |

| | | |
|---|---|---|
| Buffer N | Sld3/7 Prephosphorylation | 25 mM HEPES-KOH pH 7.5, 500 mM KCl, 5 mM $Mg(OAc)_2$, 10% v/v Glycerol, 0.02% w/v NP-40, 1 mM DTT |
| Buffer O | MCM-DH Loading | 25 mM HEPES-KOH pH 7.5, 100 mM K-Glutamate, 10 mM $Mg(OAc)_2$, 1 mM ATP, 0.02% NP-40 |
| Buffer P | MCM-DH Loading | 25 mM HEPES-KOH pH 7.5, 100 mM K-Glutamate, 10 mM $Mg(OAc)_2$, 0.02% NP-40 |
| Buffer Q | MCM-DH Loading | 25 mM HEPES-KOH pH 7.5, 500 mM NaCl, 5 mM $Mg(OAc)_2$, 0.02% NP-40 |
| Buffer R | Pre-IC/CMG Salt Stability | 25 mM HEPES-KOH pH 7.5, 5 mM $Mg(OAc)_2$, 10% v/v Glycerol, 0.02% NP-40 |
| Buffer S | DNA Replication | 25 mM HEPES-KOH pH 7.6, 100 mM K-Glutamate, 10 mM $Mg(OAc)_2$, 5 mM ATP, 0.02% w/v NP-40, 2 mM DTT |
| Buffer T | Reconstitution: phospho-DH-3745 | 25 mM HEPES-KOH pH 7.5, 100 mM KOAc, 0.02% NP-40, 25 mM D-Biotin |

**Supplementary Table 1.** List of buffers and their composition.
