## Extended Data Table 1 for "Structure of the Pre-Initiation Complex Explains CMGE Biogenesis"

| | **Pre-IC Monomer**<br>EMDB-53970<br>PDB 9RHI | **Pre-IC Dimer**<br>EMDB-53971<br>PDB 9RHJ | **DH-Sld3-MBD**<br>EMDB-53972<br>PDB 9RHL | **DH-3745**<br>EMDB-53973<br>PDB 9RHM |
|---|---|---|---|---|
| **Data collection and processing** | | | | |
| Microscope | Titan Krios | Titan Krios | Titan Krios | Titan Krios |
| Voltage (kV) | 300 | 300 | 300 | 300 |
| Camera | Falcon IV | Falcon IV | K2 Summit | K2 Summit |
| Magnification | 130,000 x | 130,000 x | 130,000 x | 130,000 x |
| Pixel size (Å) | 0.95 | 0.95 | 1.08 | 1.08 |
| Total electron exposure ($e^{-}$/Å$^2$) | 39.0 | 39.0 | 49.8 | 49.8 |
| Exposure rate ($e^{-}$/pixel/s) | 6.48 | 6.48 | 6.80 | 6.80 |
| Number of frames collected | 31 | 31 | 32 | 32 |
| Defocus range (µm) | -2.0 to -3.0 | -2.0 to -3.0 | -1.1 to -2.5 | -1.1 to -2.5 |
| Automation software | EPU | EPU | EPU | EPU |
| Energy filter slit width (eV) | 20 | 20 | 20 | 20 |
| Micrographs collected | 59,347 | 59,347 | 105,652 | 105,652 |
| Total extracted particles | 1,572,464 | 1,572,464 | 5,610,789 | 5,610,789 |
| Final particle images | 302,350 | 302,350 | 359,025 | 72,693 |
| Symmetry imposed | C1 | C1 | C2 | C1 |
| Map resolution (global, Å) | | | | |
| FSC 0.143 (unmasked) | 4.8 | 6.6 | 3.9 | 7.9 |
| FSC 0.143 (masked) | 3.2 | 3.4 | 3.1 | 3.7 |
| **Model composition** | | | | |
| Non-hydrogen atoms | 45,962 | 91,688 | 64,411 | 68,681 |
| Protein residues | 5,581 | 11,144 | 7,872 | 8,401 |
| Ligands ($Zn^{2+}$/ADP/$Mg^{2+}$) | 5/1/1 | 10/2/2 | 10/4/4 | 10/4/4 |
| DNA residues | 62 | 120 | 106 | 106 |
| **Model refinement** | | | | |
| Initial model used (PDB code) | 7PMK | 9RHI | 7P30 | 7P30 |
| Model resolution cutoff (Å) | 0.5 | 0.5 | 0.5 | 0.5 |
| Model-Map CC ($CC_{mask}$/$CC_{box}$/ $CC_{peaks}$/$CC_{volume}$) | 0.66/0.72/0.51/0.66 | 0.67/0.76/0.50/0.66 | 0.81/0.81/0.70/0.80 | 0.77/0.79/0.65/0.76 |
| Model-Map FSC 0.5 (masked/unmasked, Å) | 3.2/3.4 | 4.6/6.3 | 3.4/3.5 | 4.1/4.3 |
| ***B* factors (Å$^2$)** | | | | |
| Average B-factors (Å$^2$) | | | | |
| Protein | 179.49 | 241.77 | 155.61 | 157.62 |
| Ligand | 181.50 | 238.67 | 163.41 | 159.29 |
| DNA | 300.40 | 370.34 | 281.84 | 298.90 |
| **RMS deviations** | | | | |
| Bond lengths (Å) | 0.002 | 0.002 | 0.005 | 0.002 |
| Bond angles (°) | 0.520 | 0.466 | 1.072 | 0.514 |
| **Validation** | | | | |
| MolProbity score | 1.57 | 1.51 | 0.77 | 1.21 |
| CaBLAM outliers | 1.23 | 1.17 | 1.44 | 1.34 |
| Clashscore | 7.99 | 7.15 | 0.90 | 4.31 |
| Poor rotamers (%) | 0.02 | 0.02 | 0.36 | 0.00 |
| C-beta deviations (%) | 0.00 | 0.00 | 0.05 | 0.00 |
| **Ramachandran plot** | | | | |
| Favored (%) | 97.29 | 97.44 | 98.55 | 98.09 |
| Allowed (%) | 2.71 | 2.56 | 1.45 | 1.91 |
| Disallowed (%) | 0.00 | 0.00 | 0.00 | 0.00 |
