## Extended Data Table 2 for "Structure of the Pre-Initiation Complex Explains CMGE Biogenesis"

| | sCMGE with Sld2 and RPA EMDB-56898 PDB 28VY | sCMGE without Sld2 and RPA EMDB-56897 |
|---|---|---|
| **Data collection and processing** | | |
| Microscope | Titan Krios | Titan Krios |
| Voltage (kV) | 300 | 300 |
| Camera | Falcon IV | Falcon IV |
| Magnification | 130,000 x | 130,000 x |
| Pixel size (Å) | 0.95 | 0.95 |
| Total electron exposure ($e^{-}/Å^2$) | 38.6 | 42.0 |
| Exposure rate ($e^{-}$/pixel/s) | 6.40 | 5.57 |
| Number of frames collected | 31 | 29 |
| Defocus range (μm) | -2.0 to -2.9 | -1.4 to -2.4 |
| Automation software | EPU | EPU |
| Energy filter slit width (eV) | 20 | 20 |
| Micrographs collected | 50,060 | 70,337 |
| Total extracted particles | 4,827,716 | 2,390,783 |
| Final particle images | 1,125,095 | 72,370 |
| Symmetry imposed | C1 | C1 |
| Map resolution (global, Å) | | |
| FSC 0.143 (unmasked) | 3.4 | 4.3 |
| FSC 0.143 (masked) | 2.7 | 3.4 |
| **Model composition** | | |
| Non-hydrogen atoms | 54,130 | |
| Protein residues | 6649 | |
| Ligands ($Zn^{2+}$/ATP/ADP/$Mg^{2+}$) | 7/5/1/6 | |
| DNA residues | 35 | |
| **Model refinement** | | |
| Initial model used (PDB code) | 7PMK | |
| Model resolution cutoff (Å) | 0.5 | |
| Model-Map CC ($CC_{mask}$/$CC_{box}$/ $CC_{peaks}$/$CC_{volume}$) | 0.89/0.89/0.81/0.89 | |
| Model-Map FSC 0.143 (masked/unmasked, Å) | 2.7/2.7 | |
| ***B* factors ($Å^2$)** | | |
| Average B-factors ($Å^2$) | | |
| Protein | 158.91 | |
| Ligand | 151.93 | |
| DNA | 321.51 | |
| **RMS deviations** | | |
| Bond lengths (Å) | 0.003 | |
| Bond angles (°) | 0.506 | |
| **Validation** | | |
| MolProbity score | 1.21 | |
| CaBLAM outliers | 0.90 | |
| Clashscore | 4.27 | |
| Poor rotamers (%) | 0.00 | |
| C-beta deviations | 0.00 | |
| **Ramachandran plot** | | |
| Favored (%) | 98.04 | |
| Allowed (%) | 1.96 | |
| Disallowed (%) | 0.00 | |
